# A tissue injury repair pathway distinct but parallel to host pathogen defense

**DOI:** 10.1101/2022.10.18.509515

**Authors:** Siqi Liu, Yun Ha Hur, Xin Cai, Qian Cong, Yihao Yang, Chiwei Xu, Angelina M. Bilate, Kevin Andrew Uy Gonzales, Christopher J. Cowley, Brian Hurwitz, Ji-Dung Luo, Tiffany Tseng, Shiri Gur-Cohen, Megan Sribour, Tatiana Omelchenko, John Levorse, Hilda Amalia Pasolli, Craig B. Thompson, Daniel Mucida, Elaine Fuchs

## Abstract

Pathogen infection and tissue injury are universal insults that disrupt homeostasis. Innate immunity senses microbial infections and induces interferons (IFNs) to activate resistance mechanisms. Applying unbiased phylogenetic analysis, we show that interleukin-24 (IL24) is among the closest evolutionary homologs to the IFN family and shares a common ancestral origin. However, in contrast to IFNs, IL24 induction occurs specifically in barrier epithelial progenitors after injury and is independent of microbiome or adaptive immunity. Surprisingly, *Il24* ablation impedes not only epidermal proliferation and re-epithelialization, but also capillary and fibroblast regeneration within the dermal wound bed. Conversely, ectopic *Il*24 induction in homeostatic epidermis triggers global epithelial-mesenchymal tissue repair responses. Mechanistically, sustained *Il24* expression depends upon both IL24 receptor/STAT3 signaling and also hypoxia-stabilized HIF1α, which converge following injury. Thus, parallel to the IFN-mediated innate immune sensing of pathogens to resolve infections, epithelial stem cells sense injury signals to orchestrate IL24-mediated tissue repair.

## INTRODUCTION

Maintaining homeostasis is a conserved hallmark of biological systems, from unicellular organisms to mammals, and is exemplified by our ability to sense and resolve disruptions such as pathogen infection and tissue injury. Unlike specialized adaptive immunity, innate immunity has evolved to sense common pathogenic molecular signatures to fight infections (*1*). Barrier epithelial tissues of the skin, lung and intestine are the first line of defense against the external environment. Upon infection, these epithelia sense pathogen-associated molecular patterns (*PAMPs*) such as ‘non-self’ bacterial DNA or viral RNA. In so doing, they activate interferon response transcription factors (IRFs), which in turn promote induction and secretion of type-I and III interferons (IFNs). Upon receptor engagement, IFNs cause activation of Janus tyrosine kinases (JAKs) that phosphorylate and robustly activate transcription factors STAT1/2 to orchestrate cell, tissue and organismal level pathogen defense to resist and eliminate pathogens, allowing the host to return to homeostasis (*2*).

Injury is another acute tissue insult that multicellular organisms encounter throughout life. Wounds damage epithelial tissue and trigger a series of ordered and conserved host responses aimed at tissue repair and restoration of homeostasis (*3, 4*). Following injury, hemostasis initiates scab formation, while neutrophils and other immune cells enter the damaged tissue to launch inflammation and optimize debris clearance (Figure 1A). Subsequently, skin heals through a process of re-epithelialization and dermal remodeling. This includes tightly coordinated migration and proliferation of epidermal stem cells (EpdSCs) (*5-9*) and dermal cells such as fibroblasts, and angiogenesis in order to restore skin homeostasis (*4, 10-14*) (Figure 1A).

**Figure 1.**
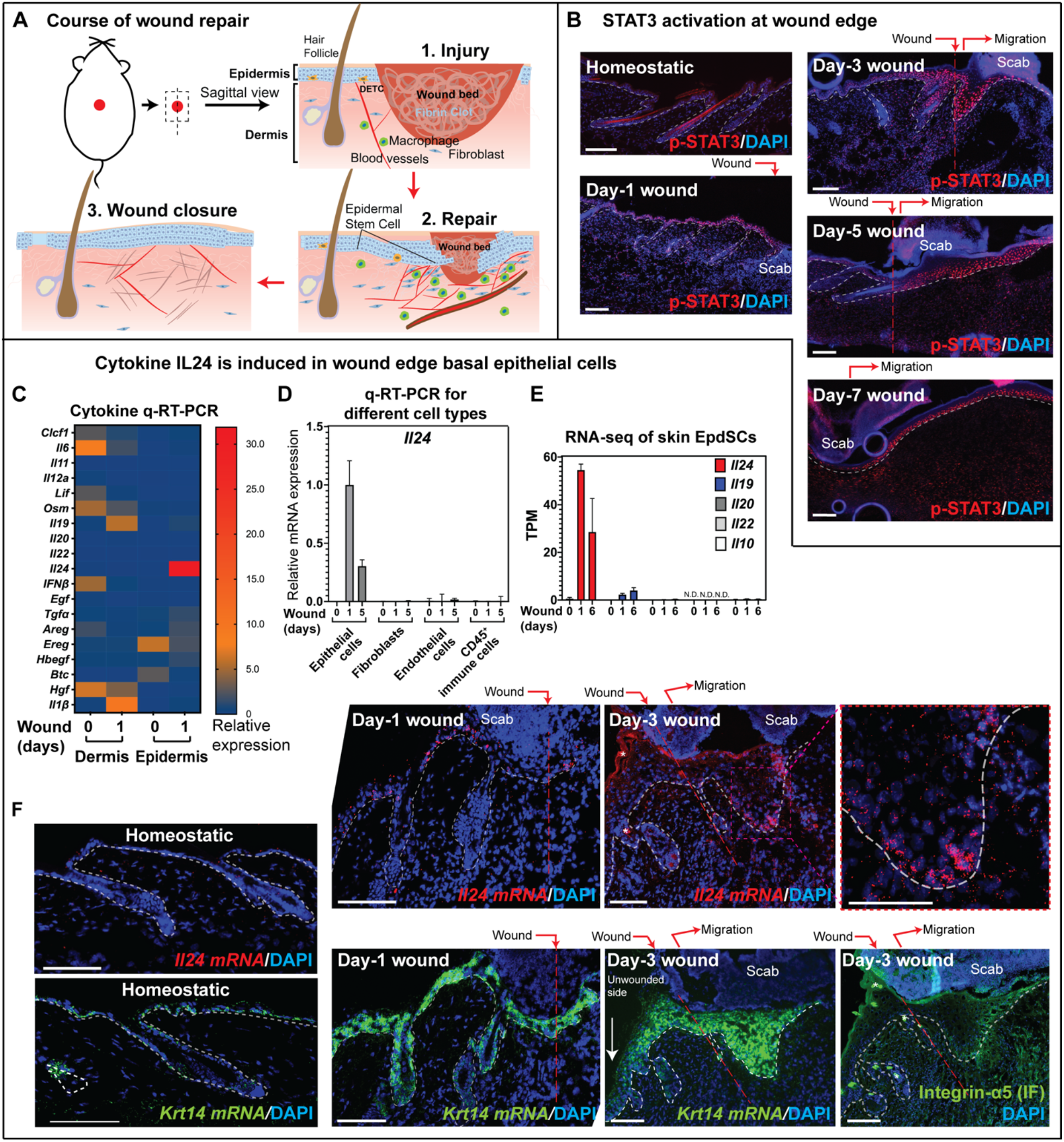
IL24 is specifically produced by epithelial stem cells near the wound site. (A) Schematic of the wound repair process in mouse skin. (B) Fluorescence microscopy images of the sagittal sections of homeostatic skin, and Day-1, 3, 5, and 7 wounds immunolabeled for phosphorylated STAT3 at Tyr705 (p-STAT3) and stained with DAPI to label nuclei. Wound site and direction of epidermal migration are indicated. Scale bars: 100 µm. [n=5 mice] (C) q-RT-PCR analysis of putative STAT3 targeting cytokines in epidermis and dermis of homeostatic skin (Day-0) and Day-1 wound. *Il1β* is known to be induced upon wounding and served as a positive control (*59*). All values were normalized to *Ppib*. [n=3 mice] (D) q-RT-PCR analysis of *Il24* mRNA expression in FACS-purified cell populations isolated from homeostatic skin (Day-0), and Day-1 and 5 wounds. [n=3 mice]* (E) IL10 cytokine family expression from RNA-seq performed on EpdSCs that were FACS-purified from homeostatic skin (Day-0), and Day-1 and 6 wounds. TPM: Transcripts per kilobase million. [n=2 mice]* (F) PLISH (proximity ligation based *in situ* hybridization) images of sagittal sections of homeostatic skin, and Day-1, 3, 5, and 7 wounds, probed for *Il24* and *Krt14* mRNA expression, and for DAPI to label nuclei. Fluorescence microscopy of anti-integrin-α5 and *Il24* PLISH in Day-3 wounds was performed on serial sections. The red-boxed region of the Day-3 *Il24* PLISH image was magnified and shown at the right to highlight the *Il24* PLISH signal in the re-epithelializing (migrating) epidermis. Asterisk (*) denotes autofluorescence in hair follicles and stratum corneum. Wound site and direction of epidermal migration are indicated. Scale bars: 100 µm; white dotted lines: epidermal-dermal border; red dotted line: angle of the incision. [n=3 mice] *The data shown in (D) and (E) are presented as mean ± SEM. N.D.; not detected.

The molecular details underlying the complex wound repair process are still unfolding. Recent studies have begun to reveal how tissue damage triggers immediate inflammatory responses in immune cells such as neutrophils and macrophages (*15-17*). How injury is sensed by the host to trigger a coordinated tissue-level repair response throughout the healing course is poorly understood especially in mammals. As a consequence, it remains largely unknown whether responses to tissue damage resemble the innate immune response to infection and if so how.

Here, we sought to identify mechanisms of how tissue damage is sensed in mammals. Exposed at the body surface, the skin epithelium is ideal to interrogate how the host senses and responds to tissue damage (Figure 1A). We identified a wound-induced signaling pathway that is independent of microbes or adaptive immunity, but functionally analogous to pathogen-induced IFN signaling in innate immunity. We show that in this parallel but molecularly distinct pathway, stem cells within the innermost (basal) epidermal layer at the wound edge sense wound-induced hypoxia as a damage signal to induce activation and signaling of interferon homolog interleukin-24 (IL24). We provide compelling evidence that in hypoxic conditions, an autocrine IL24 and IL24-receptor signaling loop is triggered which then sustains HIF1α-mediated expression of IL24 and orchestrates pro-angiogenic wound repair and restoration of tissue homeostasis.

## RESULTS

### IL24 is specifically expressed by epithelial stem cells near the wound site

We previously found that transcription factor STAT3 in wound-experiencing skin epidermal stem cells (EpdSCs) is essential to promote their crosstalk with adjacent dendritic epithelial γδT cells (DETCs) (*11*). Upon skin wounding, robust phosphorylated (active) STAT3 (p-STAT3) is induced both in the epithelial cells and dermal cells at the wound edge as early as Day-1, and throughout the wound healing course till Day-7 (Figure 1B). However, although loss of DETCs elicits modest delays in wound re-epithelialization, epithelial-specific *Stat3* ablation is much more deleterious, resulting in a striking reduction in the proliferation and migration of EpdSCs at the wound edge (*11, 18*). This vital importance of STAT3 at the nexus between tissue damage and injury repair led us to wonder whether the STAT3 signaling and its functional roles in tissue repair might be analogous to the essential roles that STAT1/2 play in pathogen resistance (*19*).

To further probe this relation, we first compiled a list of signaling factors that have been reported to activate STAT3 (*20-22*) (Table S1). We then analyzed their early activation in response to skin injury. Following a 6 mm full-thickness wound, an ∼0.5 mm skin region surrounding the wound-site was micro-dissected at Day-0 and Day-1 post-injury. Following enzymatic separation, dermal and epidermal fractions were subjected to mRNA isolation and quantitative reverse transcription and polymerase chain reaction (qPCR). Among the cytokines capable of activating STAT3, only a few exhibited a wound-induced pattern of expression. *Il24* was the most robustly induced by injury and appeared largely if not exclusively in the epidermal fraction (Figure 1C).

IL24 is a conserved mammalian gene (Table S2). It is a member of the IL10 family, which includes IL10, IL22 and the IL20 cytokine subfamily of IL19, IL20 and IL24 (*22, 23*). Although IL24 expression has been reported in different cultured cell types and in wounds *in vivo* (*24-27*), its expression, regulation and function in a natural physiological setting have remained elusive. To pinpoint the cells expressing *Il24* in skin wounds and assess IL24’s possible importance in the repair process, we performed fluorescence activated cell sorting (FACS) and purified the major cellular constituents in and within 0.5-1 mm of the wound bed at times during re-epithelialization (Figure S1A). *Il24* was transiently expressed within the EpdSC fraction (integrin-α6^hi^Sca1^hi^CD34^neg^CD45^neg^ CD31^neg^PDGFRβ^Rneg^) at the wound site (Figure 1D). Among the other IL10 family members, only *Il19* showed weak induction in the EpdSC population during the repair process (Figure 1E).

Probing deeper, we performed 10x single cell RNA-seq of skin wounds. *Il24* mRNA was detected almost exclusively within the epithelial cell cluster (*Krt14*^+^) that co-expresses high levels of basal EpdSC marker integrin-α6 (*Itga6*) and was not detected in any other cell clusters including immune cells, fibroblasts and endothelial cells (Figure S1B). In addition, we also performed bulk RNA-seq on major cell populations from skin wounds for a higher sequencing depth. Consistently, *Il24* was predominately produced in EpdSCs, a unique expression pattern not shared by other IL10 family members (Figure S1C). These observations were further supported by recently reported 10x single cell RNA-seq data from wounds (*28*).

We next combined immunofluorescence microscopy and proximity-ligation based fluorescent in-situ hybridization (PLISH) to localize *Il24* in the wound. While *Krt14* PLISH marked the epidermis of both homeostatic and wound-induced tissue, *Il24* PLISH was only detected following injury, where it appeared within a day in the EpdSCs near wound site (Figure 1F). As wound-edge EpdSCs began to migrate into the wound bed, they induced integrin-α5^+^ migration marker (*5, 8*). By Day-3, the *Il24* PLISH signal had intensified and although the wounded tissue displayed a uniform low-grade background hybridization irrespective of cellularity, the intense cellular-specific signal was concentrated within the integrin-α5^+^ basal EpdSCs at the migrating front of tissue repair (Figure 1F). This finding was corroborated by qPCR and RNA-seq of FACS-fractionated SCA1^+^ EpdSCs, which were further gated according to their level of integrin-α5 (Figures S1C and S1D). Taken together, these data pointed to the view that as yet undetermined injury signal(s) are received by nearby EpdSCs, causing them to produce IL24 specifically at the migrating epidermal front of the wound bed.

### Injury-induced IL24 signaling resembles infection-induced interferon innate immune signaling

To gain further insights into the possible significance of IL24 in this context and the possible upstream signals involved in its induction, we next performed phylogenetic analysis on all human proteins that shared both sequence and structure similarity to IL24. Among 59 homologous proteins, the IFN family is most closely related, suggesting that they share a more recent common ancestor with the IL20 subfamily compared to the other cytokines (Figure 2A, Table S3). Moreover, when we analyzed the homology of the extracellular immunoglobulin-like (Ig-like) domains of the receptors of these 59 proteins (*21*), we discovered that IFN and IL20 receptor families clustered together and shared more sequence/structure homology compared to the other cytokine receptors (Figure 2B, Table S3). IL24 can signal through heterodimeric receptor pairs IL22RA1/IL20RB or IL20RA/IL20RB (*29*), involving one long (IL22RA1 or IL20RA) and one short (IL20RB) subunit. Intriguingly, based upon their respective long vs. short heterodimeric subunits, the receptor subunits of the IFN and IL20 families also sub-clustered into two sister clades in the evolutionary tree (long vs. short subunits), suggesting that these two cytokine families likely shared a common ancestral heterodimeric receptor complex (Figure 2B). During infection, pathogen-derived signals trigger a host innate immune response for interferon production in order to clear the pathogen (*1*). Given the homology shared by IFN and IL24, and that wounds cause a skin barrier breach exposing EpdSCs to microbes, we first tested whether commensal bacteria/microbes at the skin surface are responsible for inducing *Il24* expression following injury. Notably, however, when compared with specific-pathogen-free (SPF) mice, mice that were raised under completely sterile (germ-free) conditions still showed robust induction of *Il24* in EpdSCs at the wound edge (Figure 2C). Similarly, *Il24* was still robustly induced following a wound to *Myd88*^*-/-*^*Trif*^*-/-*^ mice (Figure 2D). These mice are deficient in Toll-like receptor (TLR) signaling, which is known to mediate many microbial responses and which also plays a major role in the production of some IL10 family members (*30, 31*). Taken together, these results provided compelling evidence that distinct from the trigger of type-I IFNs by pathogens, a microbe-independent damage signal induces *Il24* at the wound edge.

**Figure 2.**
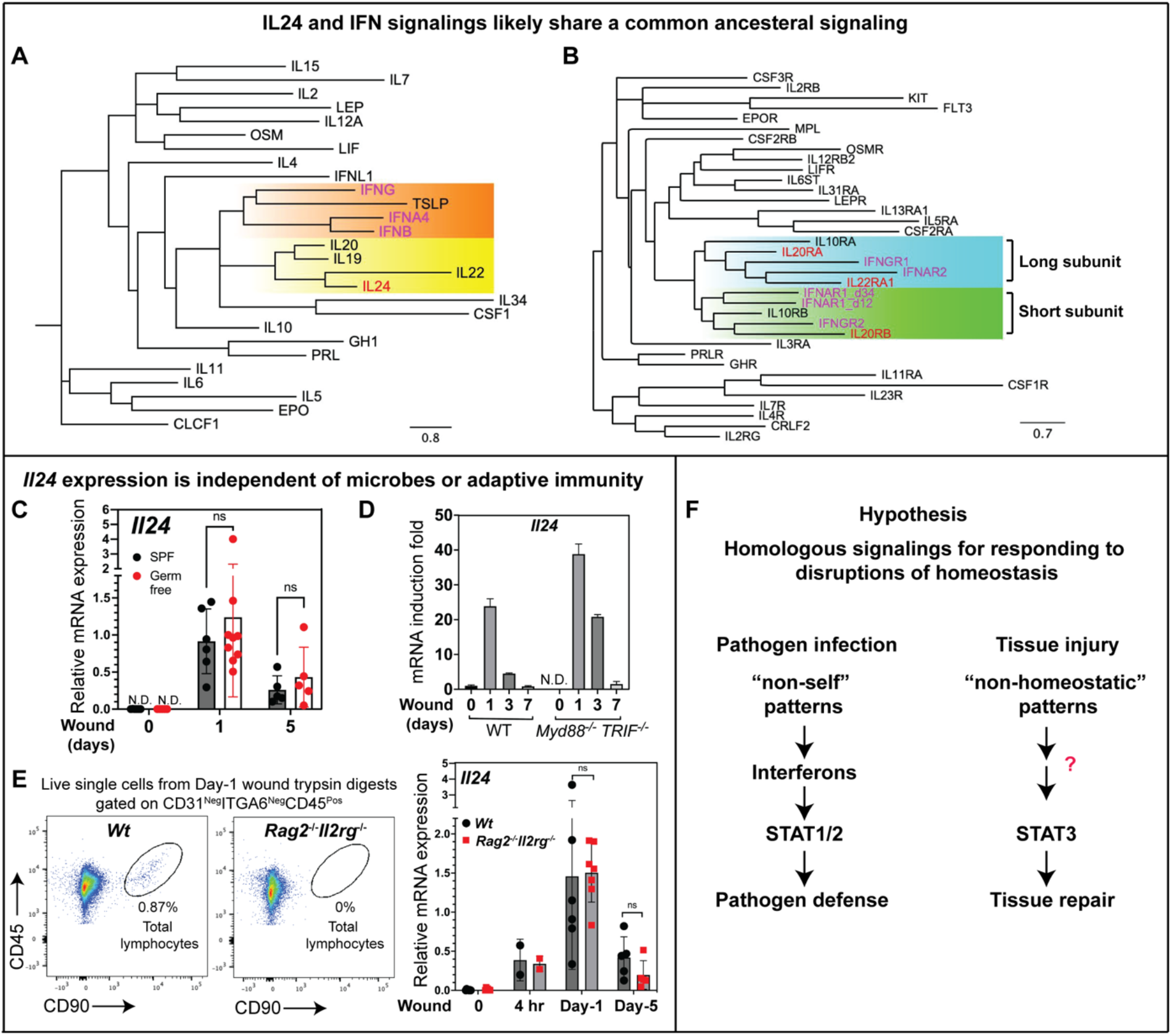
Injury-induced IL24 signaling in epidermal stem cells resembles infection-induced interferon signaling in innate immune cells. (A) Unbiased sequence and structure-based phylogenic analysis of human cytokines that share homology (see STAR method for detailed approaches). Family clades for the IL20 subfamily and IFN are highlighted in yellow and orange, respectively. (B) Sequence and structure-based phylogenic analysis of receptors for cytokines identified in (A). The clades for the long receptor subunits of the IL20 subfamily and IFN are highlighted in blue, and the clades for the short receptor subunits of these families are in green. (C) q-RT-PCR analysis of *Il24* mRNA expression in EpdSCs that were FACS-purified from homeostatic skin (Day-0), and Day-1 and 5 wounds from specific-pathogen-free (SPF) vs. Germ-free (GF) C57BL/6J WT mice. [For each time point, SPF: n=5-6 mice, GF: n=5-9 mice; dots indicate data from individual mice.]* (D) q-RT-PCR analysis of *Il24* mRNA expression in epidermis micro-dissected from homeostatic skin (Day-0), and Day-1, 3, and 7 wounds from WT vs. *Myd88*^*-/-*^*Trif*^*-/-*^ mice. Note that *Myd88*^*-/-*^*Trif*^*-/-*^ mice are compromised in their ability to respond to pathogens. [n=3 mice for each genotype; representative of three independent experiments]* (E) q-RT-PCR analysis of *Il24* mRNA expression in EpdSCs FACS-purified from homeostatic skin (Day-0), and 4 hr, Day-1, and 5 wounds from WT vs. *Rag2/IL2rg* DKO mice. Note that *Rag2/IL2rg* DKO mice lack all functional lymphocytes. [n=5-7 mice for each genotype; dots indicate data from individual mice]* (F) Diagram depicting our central hypothesis that parallel but distinct molecular signaling pathways are used for responding to and resolving homeostatic disruptions such as pathogen infection and tissue injury. The steps tackled in the current study are highlighted by a red question mark. *The data shown in (C), (D), and (E) are presented as mean ± SEM. Statistical significance was determined using Student’s t-tests; ns; not significant; N.D.; not detected.

Type-I IFNs are induced by the activation of innate immune pathways, whereas type-II IFN (IFN-γ) is predominantly induced in lymphocytes (*32*). Recent studies show that adaptive immune cells involving regulatory T cells and Rorγt^+^ T cells are important for wound repair (*9, 33*). We thus examined whether the adaptive immune system might be responsible for the induction of IL24 expression in wound edge EpdSCs. However, when compared against wild-type (WT) mice, *Rag2/IL2rg* double knockout (DKO) mice which lack functional lymphocytes altogether, still showed robust temporal *Il24* induction in EpdSCs isolated from the wound site at 4 hours, 1 day and 5 days following injury (Figure 2E) (*34, 35*). These indicate that the upstream damage signal(s) that induces *Il24* at the wound site is independent of adaptive immune cells.

Based upon the collective evidence above, we hypothesized that analogous to the sensing of pathogen-derived “non-self” patterns that prompt somatic cells to activate type I IFN-receptor-STAT1/2 signaling in defense against microbial infections, injury-induced signals that don’t exist in homeostatic conditions (“non-homeostatic” patterns) may be sensed directly by EpdSCs at the wound edge to trigger the activation of IL24-receptor-STAT3 signaling to initiate tissue damage-mediated repair (Figure 2F).

### p-STAT3 and epithelial proliferation rely upon IL24 in wound repair

To interrogate the physiological significance of IL24 in wound repair, we engineered *Il24*^-/-^ mice by directly injecting *Il24* guide RNA and CAS9 protein into fertilized embryos. Two independent CRISPR-Cas9-generated *Il24*^*-/-*^ lines were generated that harbored -4 and -23 nucleotide deletions, respectively, within exon 2, causing loss-of-function frame-shifts and premature stop codons (Figures 3A and S2A).

**Figure 3.**
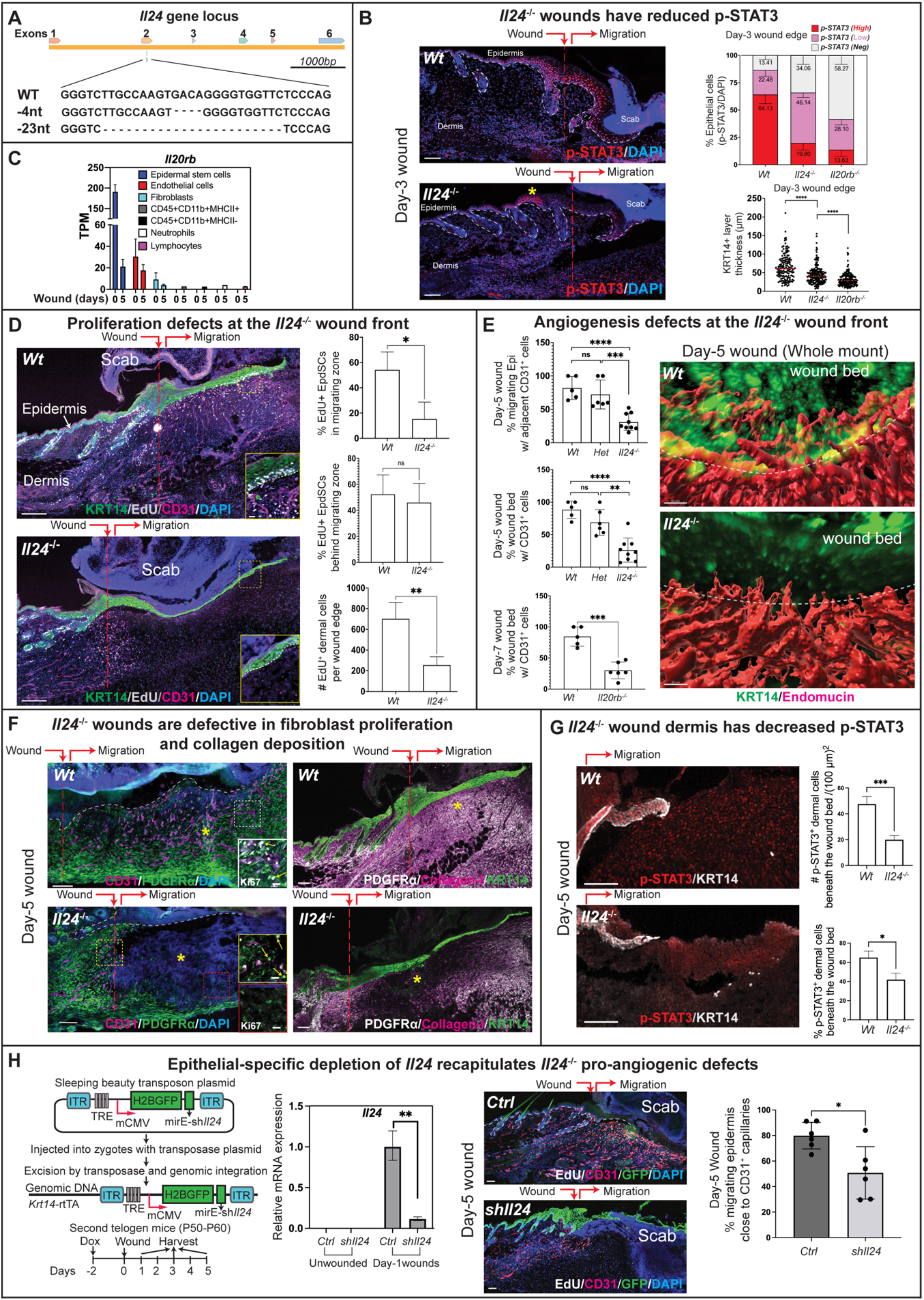
Epithelial-expressed IL24 coordinates dermal repair and re-epithelialization. (A) Schematic showing the generation of two C57BL/6J *Il24*^-/-^ mouse strains by CRISPR-Cas9-mediated frameshift deletions within the genomic locus of *Il24* exon 2. Impairments of wound repair were indistinguishable between the strains, and two loss-of-*Il24-*function strains were used interchangeably for the experiments. (B) Fluorescence microscopy images of the sagittal sections of Day-3 wounds from wild-type (*Wt*) vs. *Il24* null mice immunolabeled for p-STAT3 and stained with DAPI to label nuclei. Wound site and direction of epidermal migration are indicated. Note that p-STAT3 is still seen in *Il24* null wounded epidermis (asterisk). Scale bar: 100 µm. Graphs show quantifications of the percentage of epithelial cells expressing different levels of p-STAT3 (upper), and the thickness of KRT14^+^ layer (lower). [n=5 mice for each genotype]* (C) *Il20rb* expression from RNA-seq performed on FACS-purified cell populations from homeostatic skin (Day-0) and Day-5 wounds. TPM: Transcripts per kilobase million. [n=5 mice]* (D) Fluorescence microscopy images of the sagittal sections of Day-5 wounds immunolabeled for keratin 14 (KRT14, epidermis), EdU (proliferation), CD31 (endothelial cells), and stained with DAPI to label nuclei. Wound site and direction of epidermal migration are indicated. The boxed regions are magnified in insets to better visualize EdU incorporation of S-phase cells. Scale bar: 100 µm. Graphs show quantifications of the percentage of EdU^+^ cells in epidermis and dermis. For epidermis, quantifications were performed separately for the cells in the migrating zone (to the right of the wound site) and behind the migrating zone (to the left of the wound site). [n=5 mice for each genotype]* (E) **Left**: Quantifications of the percentages of migrating epidermis displaying adjacent CD31^+^ endothelial cells (top) and the percentages of the wound beds at Day-5 and 7 post wounding that were repopulated with sprouting blood vessels (CD31^+^ cells) (middle and bottom). Mouse genotypes are as indicated. [for *Il24* genotype quantifications, *Wt*: n=5 mice, *Il24* Het: 6 mice, *Il24*^-/-^: 9 mice, One-Way ANOVA, Tukey’s multiple comparisons test; for *Il20rb* genotype quantification, *Wt*: n=5 mice, *Il20rb*^-/-^: n=6 mice, two-tailed unpaired *t*-test]* **Right**: Images of whole-mount immunofluorescence microscopy and 3D image reconstruction performed on Day-5 wounds from *Wt* vs. *Il24* null mice. Immunolabeling was for KRT14 (epidermis) and Endomucin (blood vessels). (F) Fluorescence microscopy images of sagittal sections of Day-5 wounds from *Wt* vs. *Il24* null skins. Sections were immunolabeled for CD31, PDGFRα, and DAPI (left), or for PDGFRα, Collagen-I, and KRT14 (right). Asterisk (*) denotes a paucity of fibroblasts (PDGFRα^+^) and their deposition of Collagen-I ECM in the dermis of *Il24*^-/-^ skin. Wound site and direction of epidermal migration are indicated. The boxed region magnified in the color-coded insets shows additional Ki67 immunolabeling. Yellow arrows denote Ki67^+^ proliferating fibroblasts (Ki67^+^PDGFR^+^). Scale bars: 100 µm (20 µm in magnified images). [n=5 for each genotype] (G) Fluorescence microscopy images of the sagittal sections of Day-5 wounds immunolabeled for p-STAT3 and KRT14. Wound site and direction of epidermal migration are indicated. The boxed regions are magnified in insets to better visualize the cells expressing p-STAT3 in dermis. Scale bar: 100 µm. The percentage and the number of dermal cells beneath the wound bed that express p-STAT3 are quantified. [n=3 mice for each genotype]* (H) **Left**: Schematic of the use of a Sleeping Beauty system to generate epithelial-specific *Il24* mRNA knock-down mice. **Middle**: q-RT-PCR analysis of *Il24* mRNA expression in EpdSCs that were FACS-purified from homeostatic skin (Day-0), and Day-1 wound from control (Ctrl) vs. sh*Il24* mice. [n=5 mice for each genotype]* **Right**: Fluorescence microscopy images of the sagittal sections of Day-5 wounds from control (Ctrl) vs. sh*Il24* mice immunolabeled for EdU, CD31, Krt14, and stained with DAPI to label nuclei. Wound site and direction of epidermal migration are indicated. Scale bars: 100 µm. The percentage of migrating epidermis adjacent to CD31^+^ capillaries is quantified. [n=6 mice for each genotype]* *The data shown in (B), (C), (D), (E), (G), and (H) are presented as mean ± SEM. Statistical significance was determined using Student’s t-tests in Figures (B), (D), (G), and (H), and using One-Way ANOVA, Tukey’s multiple comparisons test and two-tailed unpaired *t* test in Figure (E); ****; p < 0.0001, ***; p < 0.001, **; p < 0.01, *; p < 0.05, and ns; not significant.

Adult *Il24*^*-/-*^ mice were healthy, fertile, and indistinguishable from WT littermates at baseline. Upon challenge, however, the wounded *Il24*^*-/-*^ epidermis displayed a markedly reduced ability to activate STAT3 specifically near the wound edge where IL24 was normally expressed by the migrating epidermal cells (Figures 3B and S2B). In marked contrast, ablation of IL6 showed little effect on p-STAT3 in wound-induced skin (Figure S2C), despite IL6 being oft-considered the major STAT3-activating cytokine in the skin (*36*). Further consistent with the reduction in p-STAT3 in *Il24*^*-/-*^ epidermis, the thickness of KRT14^+^ layers at the wound edge was markedly reduced compared to WT wounded skin (Figure 3B, quantifications), suggesting that IL24 acts directly on wound edge epithelium to sustain p-STAT3 in the epithelial tongue and promote repair.

Deletion of the pan subunit for IL24 receptor signaling resulted in even more pronounced defects in p-STAT3 and re-epithelialization (Comparative quantifications in Figure 3B; see images and more quantifications in Figures S2D and S2E). The accentuated phenotype in *Il20rb*^*-/-*^ over *Il24*^*-/-*^ wounded skins was likely attributable to redundancy between IL24 and IL19, as IL19 is known to signal through the IL24 receptor (*37*). In this regard, *Il19* was also induced by wounding, albeit at lower levels than *Il24* (Figures 1C and 1E). Notably, RNA-seq analysis confirmed that the shared IL24/IL19 receptor subunit IL20RB as well as the other two co-receptors are highly expressed in EpdSCs, indicative of the importance of epithelial IL24 and IL19 signaling in wound edge STAT3 activation and re-epithelialization (Figures 3C and S2F).

### Epithelial-expressed IL24 coordinates dermal repair and re-epithelialization

IL24 receptor was robustly expressed in the epidermis, and consistent with autocrine IL24 action, *Il24* deficiency profoundly impaired cell proliferation within the epithelial tongue of the migrating zone of the wound bed (Figures 3C and 3D; quantifications at right). Migration itself still proceeded, a process that can occur independently of proliferation (*6*).

Interesting, although IL24 receptor expression was much higher in the epidermal than in any dermal cell population (Figures 3C and S2F), marked proliferation defects were nevertheless seen in the dermis of *Il24*^*-/-*^ wounded skin (Figures 3D and S3A). Seeking the source of these dermal defects, we first co-immunolabeled for markers of proliferation and endothelial cells (CD31 and Endomucin), where IL24 receptor expression was appreciable. Notably, in the absence of IL24, a striking impairment arose in the sprouting of regenerating blood capillaries that normally account for about half of the proliferating dermal cells in Day-5 post-injury skin (Figure 3D and Figures S3B-S3D). Consistently, a recent-developed clearing method (*38*) in conjunction with whole-mount immunofluorescence and 3D image reconstruction of Day-5 wounded skin revealed a marked paucity of dermal blood vessel angiogenesis that normally closely associates with the overlying migrating epithelial tongue (Figures 3E and S3E). Quantifications revealed that ∼70% of the epidermis that had migrated into the wound bed of *Il24*^*-/-*^ skin lacked underlying vascular support (Figure 3E, top and middle quantifications). Without the support of blood vessels in the proximity, these migrating EpdSCs were also not proliferating (Figure 3D, insets). Consistent with the importance of IL24-IL24 receptor signaling, *Il20rb*^*-/-*^ mice exhibited a similar paucity of proliferating blood vessels migrating into the wound bed, a defect still evident even at Day-7 after wounding (Figure 3E bottom left panel; Figure S3F).

The remaining proliferating dermal cells in WT Day-5 wounds were mostly PDGFRα^+^ fibroblasts, but these too were largely absent in the *Il24*^*-/-*^ Day-5 wound bed (Figures 3F and S3C quantification). Consistently, the *Il24*^*-/-*^ wound bed displayed a paucity of type-I collagen, an essential extracellular matrix (ECM) component secreted by mature fibroblasts and which provides the structural support for the vasculature and the overlying epidermis. Finally, non-proliferating MHCII^+^ immune cells including macrophages were also reduced in the *Il24*^*-/-*^ dermal wound bed, especially underneath the migrating epidermal tongue (Figure S3G).

Toluidine blue staining of semithin tissue sections corroborated the gross morphological defects in restoration of dermal cellularity in Day-5 wounded skin of *Il24*^*-/-*^ mice (Figure S4A). Ultrastructural magnifications by transmission electron microscopy (TEM) further revealed the paucity of both mature dermal fibroblasts and abundant collagen deposition beneath the migrating epithelial tongue of the *Il24*^*-/-*^ Day-5 wound bed (Figure S4B). Further evidence of an impaired dermal repair process was noted by the prevalence of fibrin (pseudo-colored in green in Figure S4B). Normally resolved soon after wounding, the presence of fibrin clots left the migrating *Il24*^*-/-*^ epithelial tongue atop a fibrin clot rather than collagen-based ECM. The residual dermal fibrin and cell debris, including red blood cells (RBC), were also in good agreement with the paucity of macrophages (*39*) (Figure S4C).

Overall, the defects observed in *Il24*^*-/-*^ skin Day-5 wounds suggested that in the absence of IL24 signaling, the wound repair process had at least partially arrested at an early stage. Although *Il24-*null wounds eventually closed by contracture and limited re-epithelialization, the loss of IL24 had a profound impact on both the epidermis and the dermis, resulting in an overall decoupling of the normal repair process. Although the nature of downstream epidermal-dermal circuitries is likely to be complex and beyond the scope of the current study, these initial changes appeared to be rooted in IL24 signaling to its receptors, since not only endothelial cells, but also fibroblasts expressed significant levels of IL24 receptors (Figures 3C and S2F). In further agreement with a direct effect of IL24 on IL24 receptor signaling in the dermis, loss of IL24 resulted in clearly diminished p-STAT3 within the dermis underlying the wound bed (Figure 3G).

Although our data showed that IL24 is predominantly produced by wound edge epithelial cells in the skin, IL24 has previously been reported in other cell types and tissues (*40-44*). The broad range of wound-related defects upon whole body loss of IL24 function mandated the need to know whether these defects originated specifically from the inability to induce IL24 in the skin epithelium following injury. To this end, we generated tetracycline inducible, skin epithelial-specific *Il24* mRNA knock-down mice by directly injecting *Krt14-rtTA* fertilized mouse eggs with a sleeping beauty system including two plasmids encoding 1) transposes; 2) transposable elements including H2BGFP followed by sh*Il24* (miRE-sh*Il24*) driven by a tetracycline regulatory element (TRE) activatable by the doxycycline-sensitive transactivator rtTA (Figure 3H).

The majority of both founder and F1 offspring mice had efficient and stable integration of the transposon as indicated by the expression of H2BGFP in >90% of epidermal cells after the mice were fed with doxycycline (Dox) food. After wounding, Dox treatment efficiently silenced *Il24* mRNA induction in the sh*Il24* animals. Importantly and as we had observed in our studies with the full body knockout of *Il24*, wounds in epidermal-specific *shIl24* mice showed similar defects in the coordination of re-epithelialization and dermal angiogenesis (Figure 3H). Taken together, these data underscore the importance of skin epithelial-derived IL24, induced specifically at the wound edge, in coordinating wound repair processes of re-epithelialization, angiogenesis and collagen deposition.

### Ectopic IL24 induction in homeostatic skin epithelium elicits a wound-like response in the absence of injury

As IL24 is specifically activated following injury, we asked whether its ectopic activation might be sufficient to elicit a wound-like response in the absence of injury. A prior study in which IL24 was constitutively ectopically expressed in skin starting in embryogenesis led to neonatal lethality (*45*), emphasizing the necessity of an inducible approach to illuminate early specific effects of this cytokine on homeostatic skin and its physiological relevance to epidermal-dermal interactions. Using our powerful *in utero* lentiviral transduction method (*46*), we transduced the skin of mice genetic for an EpdSC (*Krt14*) specific, Dox-inducible transcriptional activator with lentiviral DNA containing *Il24* driven by a tetracycline regulatory enhancer (TRE) (Figure 4A). Dox administration then induced *Il24* specifically in the epidermis. Within 48 hours of IL24 induction in the basal epidermal progenitors of unwounded neonatal mice, radical changes arose in both epidermis and dermis. At the morphological level, IL24 induction robustly enhanced epidermal thickness, while in the dermis, signs of enhanced collagen deposition were seen (Figure 4B). These features were accompanied by a marked increase in proliferation both in the epidermis and the dermis (Figure 4C).

**Figure 4.**
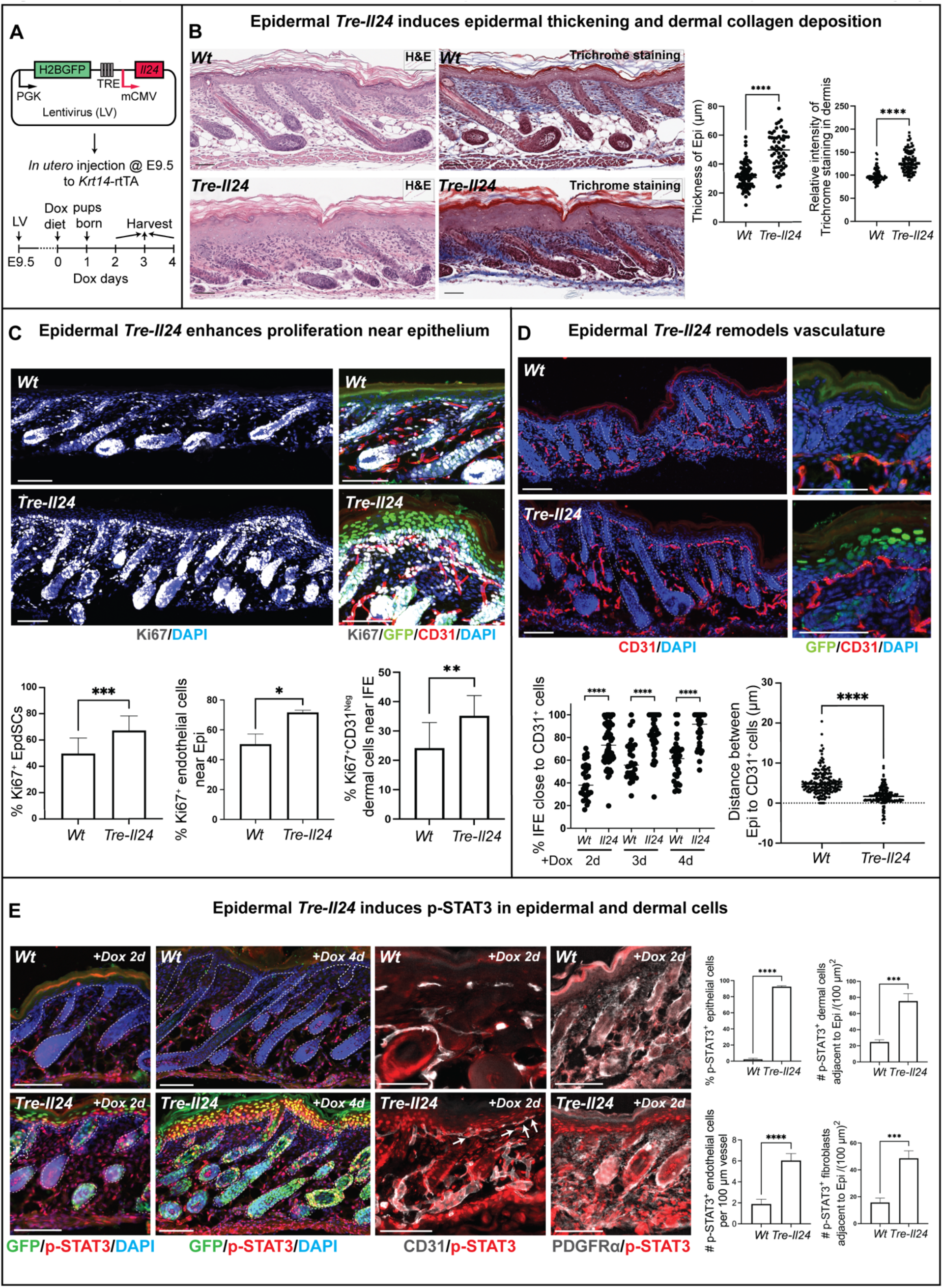
Ectopic IL24 induction in homeostatic skin epithelium elicits a wound-like response without injury. (A) Schematic of the generation of TRE-IL24 mice harboring an *Il24* transgene driven by a tetracycline regulatory enhancer (TRE). Selective targeting to skin EpdSCs was achieved by packaging the transgene in a lentivirus and *in utero* injection into the amniotic sac of E9.5 mouse embryos genetic for the *Krt14-rtTA* doxycycline inducible transcriptional activator. The lentivirus also contained a constitutively expressed *Pgk-H2BGFP* to monitor integration efficiency. Skins were harvested after the mice were fed with Dox food for 2, 3, or 4 days. (B) Images of hematoxylin and eosin (H&E) staining and Trichrome staining performed on sagittal sections of homeostatic skins from Dox-fed *Wt* and *Tre-Il24* mice. Scale bar: 100 µm. Graphs show the quantifications of the thickness of the epidermis (left), and the intensity of Trichrome staining to evaluate dermal collagen deposition (right). [n=3 mice for each genotype]* (C) Fluorescence microscopy images of the sagittal sections of homeostatic skins from *Wt* and *Tre-Il24* mice immunolabeled for Ki67, GFP, CD31, and stained with DAPI to label nuclei. Scale bar: 100 µm. Graphs show the quantifications of the percentages of proliferating (Ki67^+^) EpdSCs (left), proliferating endothelial cells (Ki67^+^CD31^+^) adjacent to the epidermis (middle), and proliferating non-endothelial dermal cells (Ki67^+^CD31^-^) adjacent to the interfollicular epidermis (right). [n=3 mice for each genotype]* (D) Fluorescence microscopy images of sagittal sections of homeostatic skins from *Wt* and *Tre-Il24* mice immunolabeled for GFP, CD31, and stained with DAPI to label nuclei. Scale bar: 100 µm; white dotted lines: epidermal-dermal border. Graphs show the quantifications of the percentage of interfollicular epidermis close to CD31^+^ endothelial cells (left), and the distance (µm) between the epidermis and CD31^+^ vasculature (right). [n=3 mice for each genotype and each time point]* (E) Fluorescence microscopy images of the sagittal sections of homeostatic skins from *Wt* and *Tre-Il24* mice stained with DAPI to label nuclei and immunolabeled for: left; GFP and p-STAT3, middle; CD31 and p-STAT3 (white arrows denote the CD31^+^p-STAT3^+^ cells), and right; PDGFRα and p-STAT3. Prior to collecting skins, mice were fed with Dox food for 2 days (+Dox 2d) or 4 days (+Dox 4 days). Scale bar: 100 µm. Graphs show the quantifications of the percentage of epithelial cells expressing p-STAT3, as well as the number of cells expressing p-STAT3 in each indicated population. [n=3 mice for each genotype and each time point]* *The data shown in (B), (C), (D), and (E) are presented as mean ± SEM. Statistical significance was determined using Student’s t-tests; ****; p < 0.0001, ***; p < 0.001, **; p < 0.01, and *; p < 0.05.

As *Il24*^-/-^ wounds experienced reduced and uncoordinated angiogenesis, we also examined the vasculature in unwounded skin that had been stimulated to ectopically express IL24. Immunofluorescence revealed that strikingly, IL24-induction in the homeostatic epidermis induced robust proliferation of CD31^+^ endothelial cells and local vascular remodeling directly beneath the EpdSC layer (Figures 4C and 4D). Quantifications provided below these images further illuminated these features.

In WT wounded skin, the strongest p-STAT3 signal was observed in the epidermal cells, which we showed also express the highest level of IL24 and IL24 receptors (Figures 1, 3C and S2F). This link appeared to be a direct one, since ectopic activation of IL24 led to a dramatic increase in epidermal p-STAT3 (Figure 4E). Consistent with the presence of lower but significant levels of IL24 receptors in endothelial cells and fibroblasts (Figure 3C and S2F), p-STAT3 also clearly rose in the dermal compartment. Indeed, co-labeling indicated that the most robust p-STAT3 dermal signals were found in the endothelial cells (CD31^+^) and fibroblasts (PDGFR?^+^) (Figure 4E), complementing our prior loss of IL24 function studies on dermal p-STAT3 (Figure 3G). Taken together, these results provided compelling evidence that in the absence of injury, epidermal-specific induction of IL24 is sufficient to elicit a tissue level wound-like response, likely through autocrine (epidermal) and paracrine (dermal) IL24 receptor signaling.

### Tissue damage associated hypoxia and HIF1α in the wounds are important for robust *Il24* expression

Given the striking role of IL24 in orchestrating epidermal-dermal dynamics during wound repair, we searched for the upstream signals that lead to *Il24* induction. Our data thus far implied that the injury signal(s) must be a “non-homeostatic” pattern that is only unleashed after wounding and which is independent of microbes or adaptive immune cells. Further corroborating this point, we also showed that this signal is independent of TNF signaling (Figure S5A), indicating that the mechanism that induces *Il24* in a physiological wound is distinct from the pathological scenario where the inhibitor of nuclear factor kappa-B kinase subunit beta (IKKβ) is deleted from skin (*26*).

In wild-type mice, epidermal proliferation during wound repair paralleled newly sprouting blood capillaries (Figure S5B), and a deficiency in dermal angiogenesis was among the most notable defects in wounded *Il24* null mice (Figures 3D and 3E). Hence, we posited that the “non-homeostatic” pattern(s) sensed by EpdSCs following injury may emanate from severed blood vessels. Turning to tissue hypoxia as a top candidate, we began by verifying that indeed the early wound bed of WT skin is hypoxic, as evidenced by robust pimonidazole labeling (Figure 5A). Correspondingly, HIF1α, the key transcription factor stabilized by hypoxia, showed robust wound-induced nuclear staining, beginning at the immediate WT wound edge following injury, and extending to the migrating (IL24-expressing) EpdSCs of the epithelial tongue during early repair (Figures 5B and S5C).

**Figure 5.**
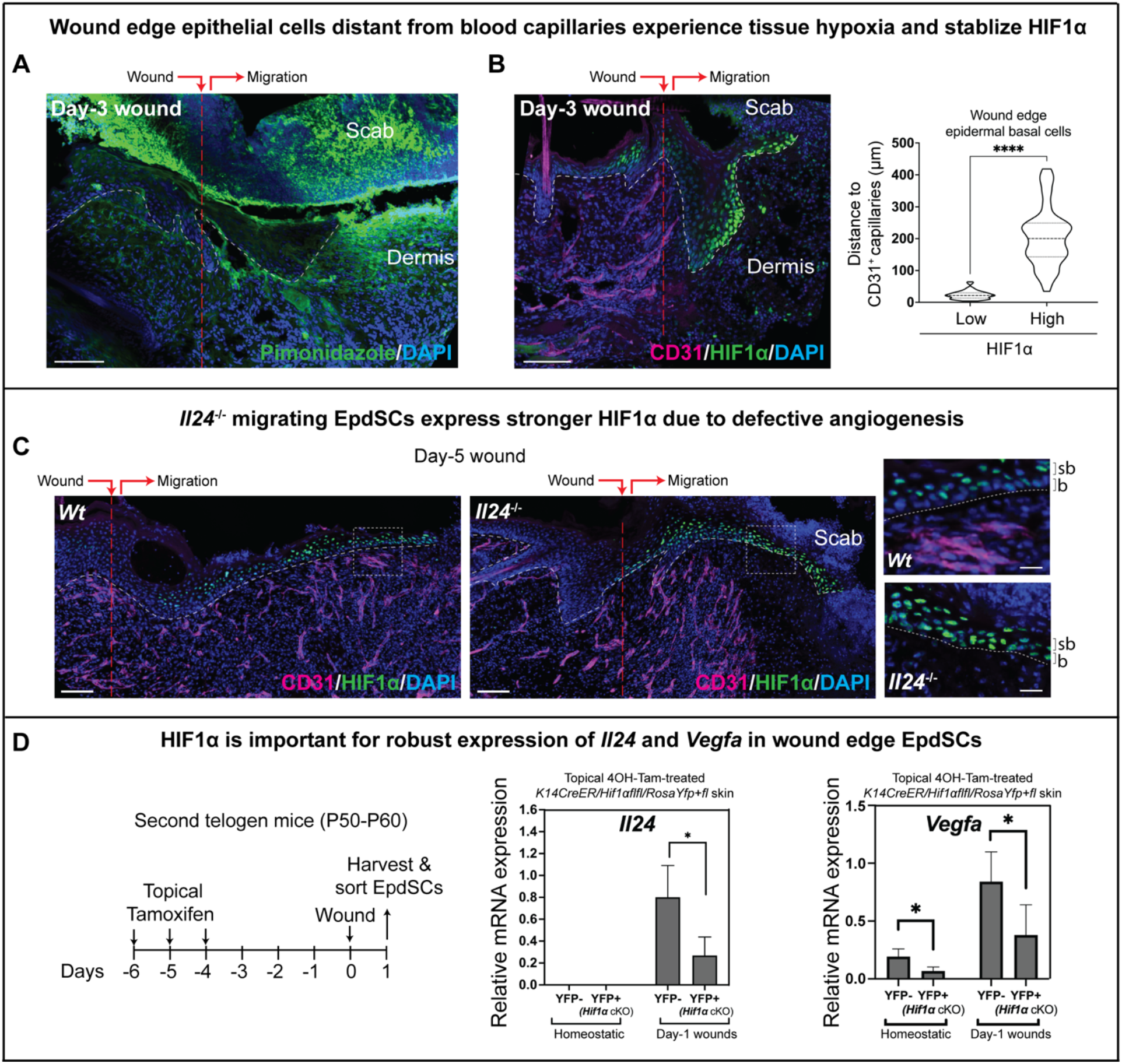
Tissue damage associated hypoxia and HIF1α in wounds are important for robust *Il24* expression. (A) Fluorescence microscopy of sagittal section of Day-3 wound harvested just after pimonidazole injection to label tissue hypoxia. DAPI was used to label nuclei. Wound site and direction of epidermal migration are indicated. Scale bars: 100 µm. [n=5 mice] (B) Fluorescence microscopy of sagittal section of Day-3 wound immunolabeled for CD31 and HIF1α and stained with DAPI to label nuclei. Wound site and direction of epidermal migration are indicated. Scale bar: 100 µm; white dotted lines: epidermal-dermal border. The distance (µm) from HIF1α^Low^ vs. HIF1α^High^ EpdSCs to the nearest CD31^+^ blood vessels is quantified. [n=5 mice, two tailed unpaired *t* test]* (C) Fluorescence microscopy of sagittal sections of Day-5 wounds from *Wt* and *Il24* null mice immunolabeled for CD31 and HIF1α and stained with DAPI to label nuclei. Wound site and direction of epidermal migration are indicated. The boxed regions of the migrating epidermal tongue are magnified in panels on the right. Scale bars: 100 µm (20 µm for magnified images); b: basal EpdSCs; sb: suprabasal epidermal cells; white dotted lines: epidermal-dermal border. [n=5 mice for each genotype] (D) Schematic of the experiment and q-RT-PCR analyses of *Il24* and *Vegfa* mRNA expression in YFP-(*Hif1α Wt*) or YFP+ (*Hif1αΔ*exon2) EpdSCs that were FACS-purified from homeostatic skin and from Day-1 wounds of *Krt14CreER; Hif1α*^*fl/fl*^; *RosaYFP*^*+/fl*^ mice treated with 4OH-Tam. Note that *Il24’s* sensitivity to hypoxia resembles that of *Vegfa*, a well-known Hif1α target gene. [n=5 mice]* *The data shown in (B) and (D) are presented as mean ± SEM. Statistical significance was determined using Student’s t-tests; ****; p < 0.0001.

Co-labeling with anti-CD31 revealed a strong correlation between the intensity of nuclear HIF1α signal in EpdSCs and their distance from blood capillaries (Figure 5B). The most robust nuclear HIF1α^+^ was always in the EpdSCs within the epithelial tongue of the wound but at least 100 µm ahead of the dermal front of regenerating (Day-3) blood capillaries. By Day-5, HIF1α had waned in basal EpdSCs concomitant with newly sprouted blood capillaries throughout the dermis, but particularly in close proximity with the overlying epidermis (Figure S5C).

By contrast, in Day-5 wounds of *Il24*^*-/-*^ mice, nuclear HIF1α remained strong in the basal EpdSCs and suprabasal epidermal cells of the migrating tongue, which lacked close contact with blood capillaries (Figure 5C). Resembling that seen in WT Day-3 wounds, the pattern of HIF1α expression in *Il24*^*-/-*^ wounds paralleled with the corresponding delay in blood capillary sprouting in the dermal wound bed and the establishment of close contact with epidermis. These data clearly placed hypoxia and HIF1α upstream from IL24, and further underscored the importance of IL24 for efficiently orchestrating wound repair.

If hypoxia plays a critical role in governing *Il24* expression, the loss of its transcriptional effector, HIF1α, might be expected to deleteriously affect wound-stimulated *Il24* induction. Indeed, this was the case, as parallel to the well-established HIF1α target gene, *Vegfa, Il24* mRNA levels plummeted when HIF1α was conditionally ablated within the epidermis prior to wounding (Figures 5D and S5D). When taken together with the importance of IL24 specifically for dermal blood capillary regeneration and EpdSC proliferation (Figures 3D and 3E), these results suggested that IL24 was induced following EpdSC-sensing of wound-generated hypoxia in order to promote revascularization and proper re-epithelialization in tissue repair.

### Critical roles for both hypoxia/HIF1α and IL24 receptor/STAT3 signaling in governing robust *Il24* expression

We next explored whether additional possible “non-homeostatic” patterns associated with blood vessel disruption could induce IL24. To this end, we established an *in vitro* primary EpdSC keratinocyte culture system and tested a panel of conditions pertinent to blood vessel disruption, including not only hypoxia, but nutrient deprivation (e.g. essential amino acids, glucose and glutamine), alternative ECM such as fibrin clots and collagen, and lactate as an abundant product from anaerobic glycolysis (Figure 6A). We also tested H_2_O_2_ as it induces oxidative stress and also has been shown to be a first signal induced by wound for immune cell recruitment (*15*).

**Figure 6.**
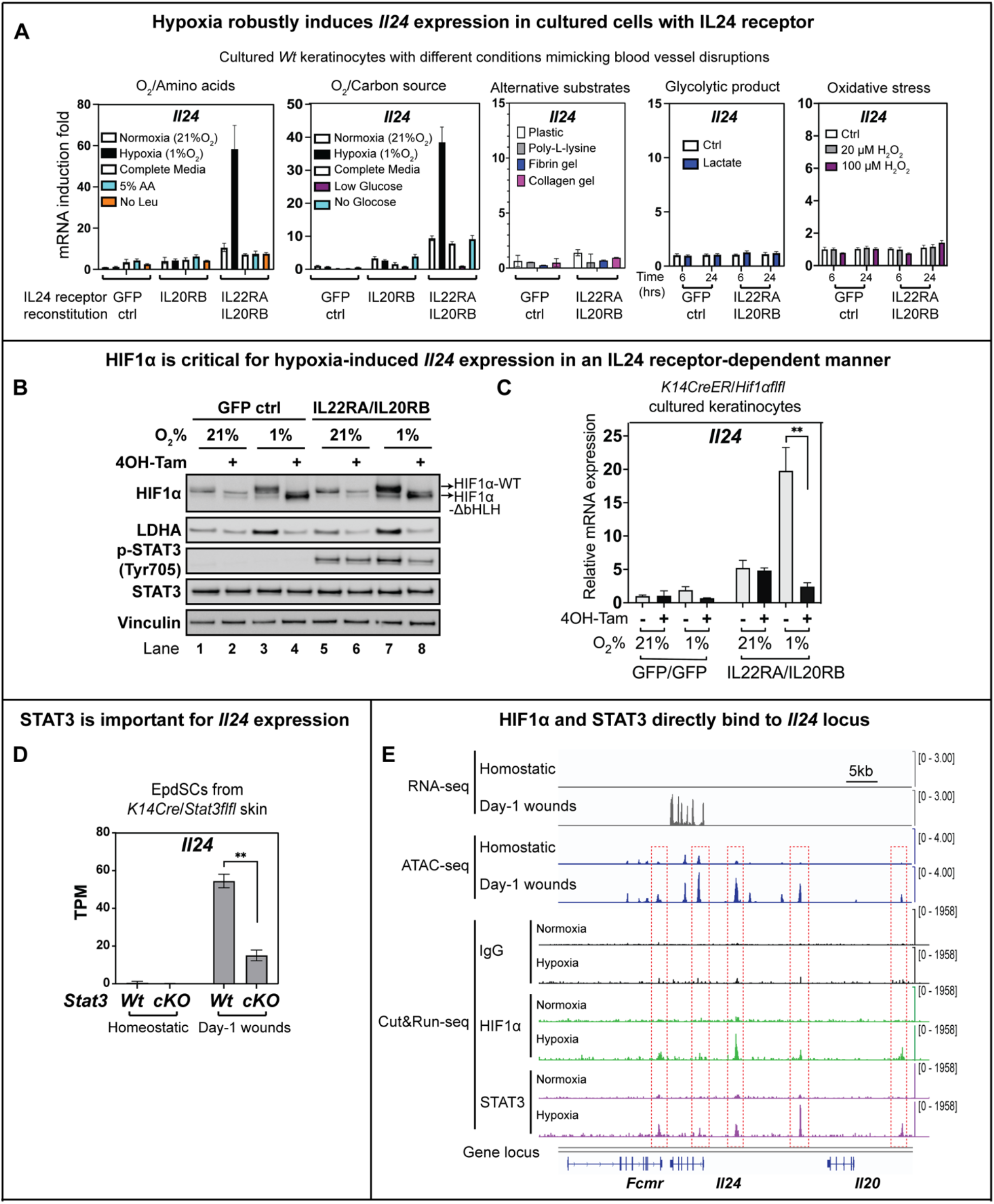
Critical roles for both hypoxia/HIF1α and STAT3 in governing robust *Il24* expression. (A) q-RT-PCR analysis of *Il24* mRNA expression in keratinocytes with GFP or IL24 receptor reconstitution cultured under different oxygen, nutrient, substrate, glycolytic product, and oxidative stress conditions for 48 hours. AA: amino acid; Leu: Leucine. Note that the native IL24 receptor, which is robustly expressed by EpdSCs in their native niche *in vivo*, is silenced under the culture conditions *in vitro*. [representative of 3 independent experiments]* (B) EpdSCs were isolated from the skin of *Krt14CreER; Hif1α*^*fl/f*^ mice, reconstituted with either GFP or IL24 receptor, and cultured in normoxic (21% O_2_) or hypoxic (1% O_2_) conditions. 4OH-Tam was used to replace the endogenous HIF1α with HIF1α lacking the bHLH DNA binding domain (*Hif1αΔ*exon2). Cells were then lysed and immunoblotted for HIF1α, LDHA (lactate dehydrogenase A; encoded by a classical hypoxia-sensitive gene), p-STAT3, STAT3, and vinculin as the loading control. [representative of 3 independent experiments] (C) q-RT-PCR analysis of *Il24* mRNA expression in the cells described in (B).* (D) *Il24* expression from RNA-seq data performed on FACS-purified EpdSCs from homeostatic skin and Day-1 wounds from *Wt* and *Krt14Cre; Stat3*^*fl/fl*^ mice treated with 4OH-Tam. TPM: Transcripts per kilobase million [WT: n=3 mice, *Stat3* null: n=3 mice]* (E) Normalized peaks of RNA-seq, ATAC-seq, and IgG, HIF1α, and STAT3 Cut&Run-seq at the *Il24* locus. Red boxes indicate the 5 chromatin regions at the *Il24* locus that opened upon wounding (ATAC), which also have both HIF1α and STAT3 binding peaks (Cut&Run). Peaks from the same experiments are indicated on the same scale. All sequencing experiments were done in duplicates. *The data shown in (C) and (D) are presented as mean ± SEM. Statistical significance was determined using Student’s t-tests; **; p < 0.01.

Unexpectedly, none of these *in vitro* conditions, including hypoxia, had a robust effect on *Il24* induction (Figure 6A). This was not because of a culture-related impairment in hypoxia-stabilized HIF1α, since traditional HIF1α targets, *Pgk1* and *Pdk1* (*47*) were markedly induced under hypoxic conditions (Figure S5E). Rather, these results suggested that *Il24* induction after injury requires not only hypoxia and HIF1α, but some additional factor(s).

Digging deeper, we learned that although the epidermal IL24 receptor, IL22RA1/IL20RB, was highly expressed *in vivo* in the EpdSCs of wounded skin, it was silenced under the *in vitro* conditions used here (Figure S5F). Upon reconstitution, IL24 receptor-positive keratinocytes displayed a robust response to hypoxia, but not to the other conditions, in eliciting *Il24* transcription (Figure 6A). Similar to the wounds, hypoxia-induced *Il24* in IL24 receptor-positive EpdSCs in culture was dependent on HIF1α (Figures 6B, 6C and S6A). These findings underscored the specific reliance upon both hypoxia/HIF1α and IL24 receptor-signaling in activating *Il24* following injury.

The existence of a positive feedback loop operative on *Il24* gene regulation was reminiscent of that seen for *Ifn* gene expression (*32*), and shed light on why following tissue damage, only EpdSCs showed robust *Il24* induction even though many skin cells experienced acute hypoxia and also stabilized HIF1α (Figures 5A and 5B). Since p-STAT3 was robustly activated in wild-type wound edge EpdSCs, and was also highly sensitive to IL24 receptor signaling in cultured EpdSCs (Figures 1B and 6B), we reasoned that like HIF1α, STAT3 might be required for optimal induction of *Il24*.

To test this hypothesis, we conditionally targeted *Stat3* in the epidermis of mice and subjected the mice to injury, as before (*11*). Notably, the induction of *Il24* in wound edge *Stat3* null EpdSCs was markedly diminished (Figure 6D). In this regard, *Il24* differed from classical HIF1α targets, e.g. *Vegfa* and *Ldha* (encoding lactate dehydrogenase A), which showed hypoxia sensitivity and functional HIF1α dependency, but were not dependent upon STAT3 for its induction (Figures 6B, S6A and S6B).

Further addressing the importance for hypoxia/HIF1α on *Il24* expression specifically, we interrogated the effects of IL17A, which is produced by wound-activated adaptive immune cells, and was recently reported to promote HIF1α stabilization after prolonged hypoxia later in the repair process (*9*). Although adaptive immune cells were dispensable for *Il24* induction *in vivo* especially early in the repair process (Figure 2E), we did find that under hypoxic conditions *in vitro*, IL17A boosted *Il24* expression (Figure S6C). These data further corroborated the importance of hypoxia for robust *Il24* expression.

Finally, to ascertain whether the *Il24* gene is a direct target of HIF1α and STAT3, and whether it is activated specifically upon wound-induced hypoxia, we subjected our purified EpdSCs from homeostatic and wounded skin (Figure S1A) to RNA-sequencing to assess *Il24* transcription and to ATAC-sequencing (Assay for Transposase-Accessible Chromatin using sequencing) (*48*) to assess chromatin accessibility at the *Il24* locus. These data are shown in Figure 6E. *Il24* transcription was robustly induced in wound edge EpdSCs. Within the *Il24* genomic locus region, several ATAC-peaks were also strongly induced in wound edge EpdSCs, suggesting that these chromatin regions were more open for transcription factor binding after wounding.

To further test this possibility, we employed Cut&Run-sequencing (*49*), which combines antibody-targeted controlled cleavage by micrococcal nuclease with massively parallel DNA sequencing to identify the binding sites of HIF1α and STAT3. We performed these experiments on primary cultured keratinocytes, where we could specifically address the sensitivity of the binding of these factors in hypoxic versus normoxic conditions and in the presence or absence of the IL24 receptors.

The data are presented in Figure 6E and S6D. Strong HIF1α and STAT3 peaks were observed within the *Il24* genomic locus. These peaks for both transcription factors occurred specifically within the ATAC-accessible chromatin regions. As a control, although *Il20* also resided within the *Il24* genomic locus, *Il20* was not expressed nor induced in the wounded EpdSC *in vivo* or *in vitro*. In addition, and consistent with our earlier findings of IL24 receptor-independent induction of *Pgk1* under hypoxia, only HIF1α and not STAT3 displayed prominent binding at the *Pgk1* locus (Figure S6D). In contrast to *Il24, Pgk1* did not show any major wound-induced expression difference when STAT3 was absent (Figure S6E).

Taken together, these data indicated that following injury, HIF1α and STAT3 are activated and directly bind to the *Il24* genomic locus, where they function together to robustly transcribe *Il24* at the wound bed. This dual dependency ensures specificity and affords fine-tuning in our response to tissue damage.

## DISCUSSION

Injury and infection are universal insults to living organisms throughout evolution. The ability to properly sense and respond to acute insults for timely resolution is essential for organismal survival. Numerous pathogen-associated molecular patterns (PAMPs) are known to stimulate innate immune signaling to resist infection (*1, 50*). How mammals sense injury at a tissue-intrinsic level is less clear (*12, 51, 52*), but is essential to advance our understanding of how initial injury sensing ultimately leads to coordinated repair and restoration of tissue homeostasis. Using skin wounding in mice as a model, we uncovered a previously elusive molecular pathway that is induced upon tissue damage, independent of microbes and the adaptive immune system but bears a striking resemblance to PAMPs-induced innate immune signaling (Figure 7).

**Figure 7.**
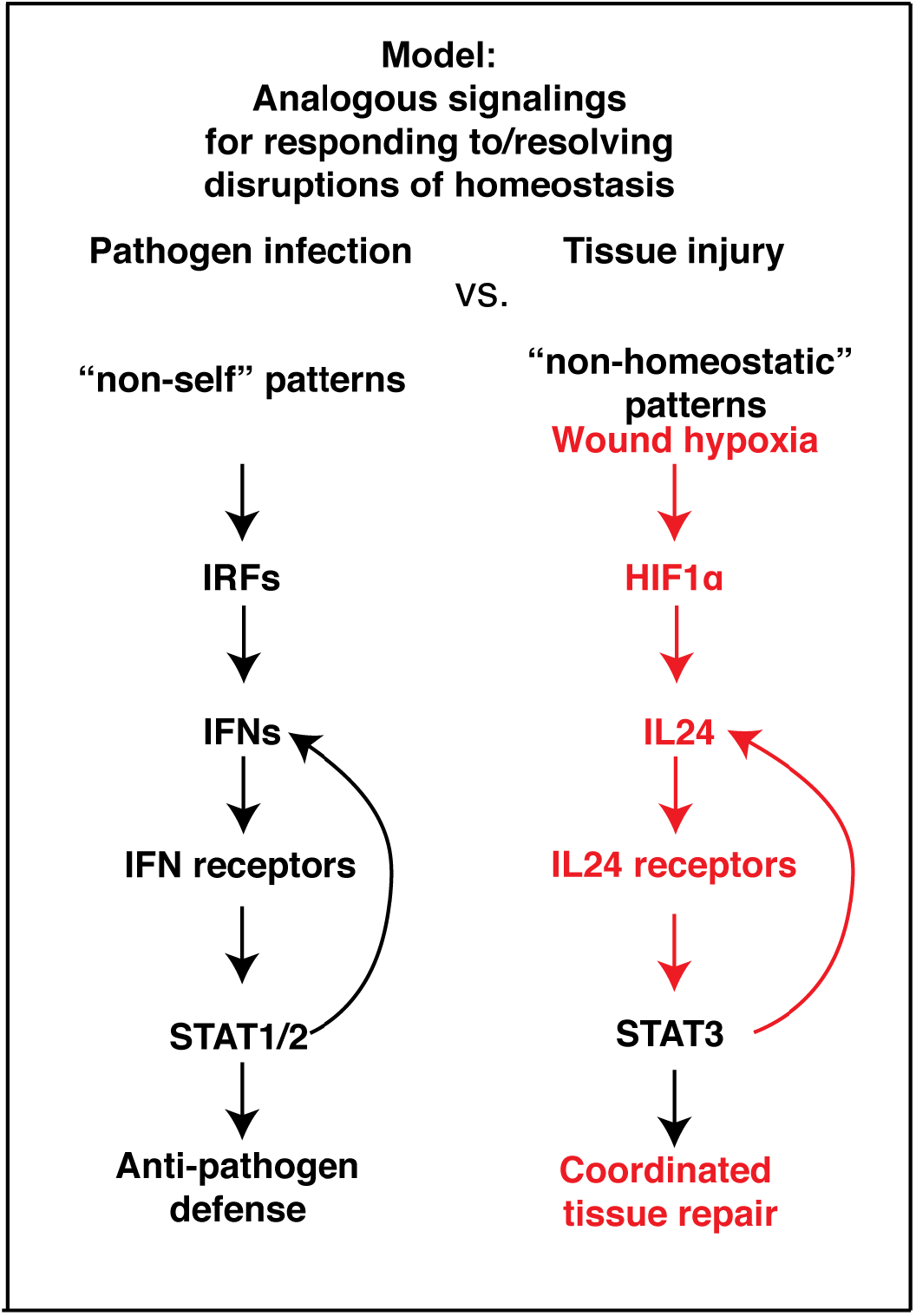
Signaling pathways activated by pathogen infection and tissue injury share striking resemblance but involve distinct molecular players. We’ve shown that upon tissue injury, EpdSCs respond to damage activated molecular patterns (DAMPs) involving hypoxia/HIF1α and IL24/p-STAT3 to launch and coordinate a global tissue repair program involving both epidermis and dermis. Specifically, EpdSCs sense wound hypoxia caused by severed blood vessels, and induce IL24 production and receptor signaling, which subsequently activate STAT3 and further fuel IL24 generation to promote a coordinated dermal repair and re-epithelialization. The mechanisms underlying wound-induced IL24 signaling in tissue repair are parallel and functionally analogous to pathogen-induced interferon signaling in pathogen defense, and the two pathways share multiple levels of homology.

At the root of this tissue damage pathway is the interferon homologue IL24, which while not expressed in homeostasis, was specifically induced by EpdSCs at the hypoxic wound region where blood vessels have been severed. The ability to sense tissue damage such as hypoxia in a microbe-independent manner distinguishes IL24 from PAMP-induced IFN signaling. However, analogous to the role of IFN in resisting pathogen infection, IL24 coordinates a pro-angiogenetic repair and proliferation program to restore tissue integrity and homeostasis. Our unbiased phylogenetic analysis exposed IFN, IL24 and their respective receptors as close evolutionary homologs derived from a common ancestral pathway. It is tempting to speculate that as we evolved, this pathway diverged to cope with the increasing diversity of homeostatic disruptions in both pathogens and injuries.

IL24 expression in skin wounds has been previously reported in rats and humans and in a variety of cell types (*25, 53*). However, by developing and employing a cohort of genetic mouse models, we were able to delve into the functions and physiological relevance of this cytokine in ways not previously possible. In doing so, we unearthed its hitherto unrecognized importance as a master coordinator of tissue repair following injury. Strikingly, we learned that IL24 is unique among the IL20 family of cytokines in that it is strongly and specifically induced by the skin epithelial stem cells and only at the edge of the wound bed. Although we found evidence for some redundancy with IL19 in the injury response, our findings showed that hypoxia is noticeably more integral to IL24 than IL19 induction.

IFN production must be tightly regulated to prevent inflammation and autoimmunity (*32, 54*). We learned that IL24 production is similarly tightly regulated and occurs only at the wound site. Although the damaged blood vessels at a wound site generates an hypoxic state, hypoxia was not sufficient for *Il24* activation, but rather also required autocrine IL24 receptor expression and STAT3 activation. Although dissecting the mechanisms underlying IL24 receptor regulation is beyond the scope of the present study, it was intriguing that receptor expression was significantly higher in epidermal stem cells than any other cell type within the skin. It was also markedly downregulated *in vitro*, suggesting that the epidermal stem cell niche is critical in poising the EpdSCs to respond to injury and activate IL24 receptor signaling.

Another surprising finding was the p-STAT3 activation in EpdSCs at the wound site is markedly more dependent on IL24/receptor signaling than IL6. Given the feedback loop that we exposed here, and the dependency of both HIF1α and p-STAT3 on *Il24* gene expression, the reliance on IL24 receptor signaling for maintaining p-STAT3 provides an interesting insight into how the epithelial tongue progresses specifically at the wound site and simultaneously coordinates dermal repair in proximity. In the end, the repair process becomes naturally autoregulated at the back-end in that as the vasculature is re-established, both the hypoxia-induced signaling and *Il24* expression are dampened.

The IL24-mediated tissue repair pathway we delineated here is likely to have significance that extends beyond skin wound repair. Indeed, barrier epithelia like the skin are susceptible to both tissue injury and microbial infections, which often occur together. While wounding causes direct physical injury, infection can lead to secondary tissue damage, the proper repair of which is essential for disease tolerance and host survival (*55, 56*). In this regard, it is intriguing that severe COVID-19 patients who experience lung damage display prominent IL24 and Amphiregulin (Areg), another protective molecule of lung damage (*57, 58*), while the colon of patients with ulcerative colitis also show IL24 expression (*43*). Our findings offer insights into these complex disease settings of infectious and inflammatory diseases, where induction of IL24 is likely triggering an injury repair mechanism similar to the one we unraveled in our study.

In closing, we show that epithelial stem cells function as major sensors of tissue injury. Through sensing injury signals such as hypoxia and autocrine IL24 receptor/p-STAT3 signaling to maximize IL24 production, IL24-stimulated stem cells not only choreograph their own proliferation and re-epithelialization to seal wounds, but also coordinate the requisite dermal repair responses that involve blood vessel sprouting and fibroblast reconstruction of the extracellular matrix. Indeed as our gain and loss of function studies show, IL24 signaling is vital in orchestrating wound repair, bringing this cytokine to the forefront as a potential new target for enhancing repair in chronic wound patients.

## ACKNOWLEDGMENTS

We thank J. Racelis, E. Wong, L. Polak, M. Nikolova, L. Hidalgo for technical support, and S. Ellis, R. Niec, Y. Miao, M. Schernthanner, H. Yang, M. Parigi, A. Gola, C. Ng, R. Yang, Y. Yu for discussions. We thank I. Matos, Y. Miao, L. Xi and T. Feinberg for experimental contributions. FACS was conducted by Rockefeller University’s Flow Cytometry Core (S. Svetlana, Director); ATAC-seq, Cut and Run-seq, and 10x single cell RNA-seq were conducted by RU’s Genomics Core (C. Zhao, Director); RNA-seq were conducted by Genomics Core Facility of the Weill Cornell Core Laboratories Center (J. Xiang, Director); all mouse work was performed in RU’s Center for Comparative Biology and under ALAAC accreditation and according to guidelines for animal care, set by the National Institutes of Health. E.F. is an investigator of the Howard Hughes Medical Institute. S. L. is a recipient of Rockefeller Women &Sciences Postdoctoral Fellowship, Jane Coffin Childs Postdoctoral Fellowship and National Institutes of Health K99 Transitional Award; Y. Hur is a recipient of AACR-Incyte Immuno-oncology Research Fellowship; C.X. is recipient of C.H.Li Memorial Fellowship, National Cancer Center Fellowship and Charles Revson Postdoctoral Fellowship; K.A.U.G. is a recipient of Cancer Research Institute Carson Family Fellowship, Bristol-Myers Squibb Fellowship, and Human Frontier Science Program Long-Term Fellowship. This study was supported by grants to E. Fuchs from the National Institutes of Health (R01-AR050542 and R01-AR27833) and the Starr Foundation, S.L. from the National Institutes of Health (5 K99 AR072780-02 SL) and from Rockefeller Robertson Therapeutic Development Funds, and C.B.T. from National Institutes of Cancer and cancer center support grant to Memorial Sloan Kettering Cancer Center (P30 CA008748).

## AUTHOR CONTRIBUTIONS

Conceptualization: S.L., E.F.; Methodology: S.L., Y.H., E.F., X.C., Q.C., C.X., A.M.B., B.H., C.J.C., K.A.U.G., Y.Y., T.T., S.G.C., M.S., HAP; Investigation: S.L., Y.H., B.H., M.S. (Crispr/CAS-mediated ablations *in vitro* and in mice); S.L., Y.H., C.X., J.L. (Sleeping beauty sh*Il24* mice); S.L., X.C., C.B.T. (hypoxia experiments *in vitro*); Q.C. (evolutionary tree analyses); S.L., C.X. (PLISH); S.L., Y.H., T.O. (Tre-IL24); S.L., C.J.C., K.A.U.G., Y.Y., J.L. (high throughput studies/analyses); S.L., H.A.P. (ultrastructural studies); S.L., S.G.C. (whole mount 3D clearance of wounds); S.L., A.M.B., D.M. (germ-free facility studies); S.L., Y.H., T.T. (all other experiments); Visualization: S.L., Y.H., E.F.; Funding acquisition: E.F., S.L; Supervision: S.L., E.F; Writing – original draft: S.L., Y.H., E.F.

## INCLUSION AND DIVERSITY STATEMENT

The authors are a team of scientists from multiracial background and with collective international training experiences. There are equal numbers of female and male authors. One or more authors of this manuscript is self-identified as a member of LGBQ+ community. We strongly support the diversity in scientific community.

## MATERIALS AND METHODS

### RESOURCE AVAILABILITY

#### Lead contact

Further information and requests for resources and reagents should be directed to and will be fulfilled by the lead contact, Elaine Fuchs.

#### Materials availability

Materials used in this study will be provided upon request and available upon publication.

#### Data and code availability

All other data in the manuscript, supplementary materials and source data are available from the corresponding author upon request.

- Bulk RNA-, 10x singe-cell RNA-, ATAC-sequencing data and Cut-and-Run sequencing data from this study have been deposited in the Gene Expression Omnibus (https://www.ncbi.nlm.nih.gov/geo/) under accession codes PRJNA731164, PRJNA885018, and PRJNA731304.
- All original code is available from the lead contact upon request.
- Any additional information required to reanalyze the data reported in this paper is available from the lead contact upon request.

### EXPERIMENTAL MODEL AND SUBJECT DETAILS

#### Animals

C57BL/6 and *B6.129×1-Gt(ROSA)26Sor*^*tm1(EYFP)Cos*^*/J* (*Rosa26-stop-lox-stop YFP*) mice were purchased from The Jackson Laboratory. *Krt14-Cre* and *Krt14-CreER* mice were previously generated in the Fuchs laboratory. *Il20rb*^-/-^ mice were obtained from Genentech, which was previously used in a skin wound healing study (*60*). *Hif1α* null mice were obtained by crossing Hif1α floxed animals from The Jackson Laboratory (Stock No: 007561) to K14-CreER/Rosa26-YFP (Fuchs Lab) animals. *Stat3* cKO mice were obtained by crossing Stat3 floxed animals from The Jackson Laboratory (Stock No:016923) to *K14-Cre/Rosa26-YFP* (Fuchs Lab) animals. *Myd88*^*-/-*^(Stock No: 009088) and *Trif*^*-/-*^ (Stock No: 005037) mice were obtained from Jackson Laboratories and crossed into *Myd88*^*-/-*^*Trif*^*-/-*^ in-house. *Rag2*^*-/-*^*Il2rg*^*-/-*^ (Stock No. 4111-F) and control wildtype C57BL/6NTac (Stock No. B6-F) females were purchased from Taconic. TNFR1/TNFR2 DKO mice were purchased from The Jackson Laboratory (Stock No. 003243). In order to generate *Il24*^*-/-*^ mice using the CRISPR-Cas9 method, we used the Alt-R CRISPR-Cas9 system from IDTdna. *Il24* gRNA (GGAGAACCACCCCTGTCACT) targeting its exon 1 was selected using guidescan (http://www.guidescan.com/). crRNA (containing *Il24* gRNA sequence), tracrRNA (IDT cat. #1072533), and recombinant Cas9 (IDT cat. #1081058) were purchased from IDTdna, and crRNA:trRNA:Cas9 RNP particles were assembled *in vitro* as described by the manufacturer and suspended in injection buffer (1 mM Tris-HCl pH 7.5, 0.1 mM EDTA) at a final RNP concentration of 0.122 µM. The mixture was then injected into the pronucleus of fertilized single-cell mouse embryos, and embryos were implanted into the oviducts of pseudo-pregnant *Krt14-rtTA* C57BL/6 female mice (*61*). For the generation of mice with inducible *Il24* loss of function specifically in skin epithelium, we used the sleeping beauty system and mir-E based shRNA method (*62*). For TRE-inducible *Il24* knockdown *in vivo*, we designed *Il24* shRNA with the algorithm from splashRNA (*63*), and cloned the shRNA with the optimal antisense sequences (TAGAATTTCTGCATCCAGGTCA) into the mir-E backbone (*64*) placed at the 3’UTR of a nucleus-localized H2B-GFP reporter driven by a TRE promoter. After validation of efficient knockdown in keratinocytes *in vitro*, the TRE-H2B-GFP-shIl24 cassette was cloned into a sleeping beauty transposon (Addgene Plasmid #108352) for injection into the zygotes of *K14rtTA* mice (*65*). The transposon plasmid was then mixed with a plasmid encoding transposase (pCMV-SB100; Addgene Plasmid #34879) in injection buffer (2.5 ng/µl transposon plasmid; 1.25 ng/µl SB100 transposase plasmid; 5 mM Tris-cl pH 7.4, and 0.1mM EDTA), and injected into the pronucleus of fertilized single-cell mouse embryos of *K14rtTA*, and embryos were implanted into the oviducts of pseudo-pregnant C57BL/6 female mice. Once the sleeping beauty mice were born, female mice and control littermates were subjected to wounding experiments, while male mice with high transduction efficiency were used as founder mice to back-cross with *Krt14-rtTA* C57BL/6 female mice to generate F1 offspring mice.

Animals were assigned randomly to experimental groups and studies were not blinded. However, age-and sex-matched, and whenever possible, littermates were used for each experiment. For the full-thickness wounding experiments, mice in the telogen phase of the hair cycle (P50-P65) were used. Mice were maintained in the Association for Assessment and Accreditation of Laboratory Animal Care-accredited animal facility of The Rockefeller University (RU), and procedures were performed with Institutional Animal Care and Use Committee (IACUC)-approved protocols. Mice of all strains were housed in an environment with controlled temperature and humidity under specific-pathogen-free conditions, on 12 hour-light:dark cycles, and fed with regular rodent’s chow or doxycycline as described. The numbers of animals used for experiments presented in each figure are indicated in the legends as *n* = *x* mice per group, per time point analyzed.

#### Primary cell cultures

Primary epidermal stem cells (EpdSCs) were maintained at 37°C in a humidified atmosphere containing 7.5% CO_2_. Cells were cultured in E-low calcium (50 μM Ca^2+^) medium made in-house from DMEM/F12 (3:1 ratio) medium supplemented with 15% chelated FBS, 5 μg/mL insulin, 5 μg/mL transferrin, 2 nM triiodothyroxine, 40 μg/mL hydrocortisone, and 10 nM cholera toxin. For *in vitro* hypoxia experiments, cells with GFP or IL24 receptor reconstitution were cultured under 21% oxygen (normoxia) or 1% oxygen (hypoxia) in DMEM/F12 supplemented as described above. For the generation of each nutrient-deprived condition, amino acid/glucose/glutamine deficient DMEM/F12 (complete deficient media) was made in-house by the MSKCC media core (dialyzed chelated FBS was used), and reconstituted with each nutrient and/or BSA, and the complete medium re-supplemented with all missing nutrients served as a control. Cells were also cultured on the plates coated with poly-L-lysine, fibrin, or collagen as indicated, according to the manufacturer’s instructions.

#### Cell lines

293TN HEK cells for lentiviral production were cultured in DMEM medium with 10% FCS (Gibco) and 1 mM sodium pyruvate, 2 mM glutamine, 100 units/mL streptomycin, and 100 mg/mL penicillin.

### METHOD DETAILS

#### Full-thickness wounding

Punch biopsies were performed on anesthetized mice in the telogen phase of the hair cycle (P50-P65) (*66*). For wounding the back skin, dorsal hairs were shaved with clippers and skin was swabbed with ethanol prior to wounding. 6 mm biopsy punches (Miltex) were used to make full-thickness wounds. After wounding, tissues were collected at 1, 3, 5 or 7 days after wounding as indicated.

#### Immunofluorescence microscopy

Mouse back skin was dissected, fixed with 4% paraformaldehyde diluted in PBS for 1-2 hours at 4°C, washed with PBS three times, incubated with 30% sucrose at 4°C overnight, and then embedded in OCT (Tissue Tek). Frozen tissue blocks were sectioned at 14 µm on a cryostat (Leica) and mounted on SuperFrost Plus slides (Fisher). The tissue sections were blocked for 1 hour at room temperature with the blocking solution (5% normal donkey serum, 0.5% bovine serum albumin, 2.5% fish gelatin, and 0.3% Triton X-100 in PBS). For endomucin staining, 2.5% normal donkey serum and 2.5% normal goat serum were used in the blocking solution instead of 5% normal donkey serum. Sections were then incubated with the indicated primary antibodies diluted in the blocking solution at 4°C overnight. For staining the tissues with an anti-p-STAT3 antibody, the sections were pretreated with ice-cold 100% ethanol prior to the blocking step. The sections were then washed three times with 0.3% Triton X-100 in PBS and incubated with secondary antibodies diluted in the blocking solution at room temperature for 1 hour. Finally, the sections were washed three times with 0.3% Triton X-100 in PBS, three times with PBS containing DAPI at a 1:3,000 dilution, and then mounted with ProLong Dimond Antifade Mountant (Thermo Fisher Scientific). EdU click-it reaction was performed according to the manufacturer’s instructions (Life Technologies) after the secondary antibody incubation and was followed by washing with PBS containing DAPI, as needed. The samples were visualized with an AxioOberver.Z1 epifluorescence microscope equipped with a Hamamatsu ORCA-ER camera and an ApoTome.2 (Carl Zeiss) slider. Tiled and stitched images of sagittal sections were collected using a 20X objective, controlled by Zen software (Carl Zeiss). Alternatively, whole wound images were captured using a BioTek Cytation 5 using a 4x air objective. ImageJ software was used to project Z-stacks and process images. The size of the images was adjusted and assembled in Adobe Illustrator. Scale bars were indicated in the figures.

#### Proximity ligation in situ hybridization

Proximity ligation in situ hybridization technology (PLISH) is performed as previously described (*67*) with slight modifications. Mouse skin samples were fixed with 4% paraformaldehyde in DEPC-treated PBS at 4°C for 1 hour, rinsed three times with DEPC-treated PBS, incubated with DEPC-treated 30% Sucrose/PBS solution for a few hours, and embedded in OCT. 10 μm tissue sections were prepared from frozen OCT blocks, pretreated with 25 μg/ml pepsin in 0.1 M HCl at 37°C for 5 minutes, and rinsed with DEPC-treated PBS. After drying at room temperature for approximately 5 minutes, tissue sections on the microscope slides are sealed with adhesive chambers (Grace Bio-Labs, GBL622514), rinsed with Hybridization Buffer (1 M NaTCA, 5 mM EDTA, 50 mM Tris pH 7.4, 0.2 mg/mL Heparin, and 0.1% LDS in DEPC-treated water), and incubated with a mixture of hybridization probes (sequences listed below, 100 nM final concentration each) in Hybridization buffer at 37°C. After a 2 hour-incubation in a humid hybridization oven, the tissue sections were rinsed four times with Hybridization Buffer, incubated with High Salt Buffer (0.25 M NaCl, 50 mM Tris, 2 mM EDTA, and 0.1% LDS in DEPC-treated water) at 37°C for 10 minutes, rinsed once with Circle Hybridization Buffer (2x SSC/20% Formamide, 0.2 mg/mL Heparin, and 0.1% LDS in DEPC treated water), and incubated with 116 nM phosphorylated Common Connector Circle (CCC) oligo and phosphorylated Variable Bridge (VB) oligo (sequences listed below) in Circle Hybridization Buffer at 37°C in a humid hybridization oven. After a 1 hour-incubation, the tissue sections were rinsed twice in Circle Hybridization Buffer, once with 1x T4 DNA ligase buffer (NEB, B0202S) in nuclease-free water (Invitrogen, AM9937), and incubated with a ligation reaction mixture (10 unit/μL T4 DNA ligase (NEB, M0202M), 1x T4 DNA ligase buffer, 0.4 μg/μL BSA, 0.4 unit/μL RNaseOUT (Invitrogen, 10-777-019), 250 mM NaCl, 0.005% Tween-20 in nuclease-free water) at 37°C in a humid hybridization oven. After a 2 hour-incubation, tissue sections were rinsed twice with Circle Hybridization Buffer, rinsed once with 1x phi29 polymerase buffer (Lucigen, NxGen kit 30221) in nuclease-free water, and incubated with a rolling-circle amplification (RCA) reaction mixture (1 unit/μL phi29 polymerase (Lucigen, NxGen kit 30221), 1x phi29 polymerase buffer, 5% Glycerol, 0.25 mM each dNTP, 0.4 μg/μL BSA, 0.4 unit/μL RNaseOUT in nuclease-free water) at 37°C in a humid hybridization oven. After overnight (∼16 hours) RCA reaction, the tissue sections were rinsed twice with Label Probe Hybridization Buffer (2x SSC/20% Formamide, 0.2 mg/mL Heparin in nuclease-free water) and incubated with 50 nM Label Probe (sequence listed below) in Label Probe Hybridization Buffer at 37°C in a humid hybridization oven for 2 hours. The labeled samples were washed twice with 0.05% Tween-20 in DEPC-treated PBS, stained with 1 μg/ml DAPI in DEPC-treated PBS, rinsed with DEPC-treated PBS, and imaged on the MIDAS microscope. The DNA oligos used for PLISH were purchased from Eurofins Genomics. The sequences (from 5’ to 3’ end) are listed below:

-CCC (5’ phosphorylated, HPLC purification): ATTCCTGACCTAACAAACATGCGTCTATAGTGGAGCCACATAATTAAACCTGGCTAT

-VB (5’ phosphorylated, HPLC purification): ACTACTCGACCTATAACCATAACGACGTAAGT

-Label Probe (5’ conjugated with Alexa Fluor 647, HPLC purification): ACTATACTACTCGACCTATA

-Design of H probes:

Il24-H1L: AGGTCAGGAATACTTACGTCGTTATGGAGGGTCCTAAAGTGAAGCCG Il24-H1R: AAAGGGCCAGTGCTCCTGCTTTATAGGTCGAGTAGTATAGCCAGGTT Il24-H2L: AGGTCAGGAATACTTACGTCGTTATGGAGGCTCAGGCAGGGGAGAAT Il24-H2R: GGTTCCAAAGAAGAAGGATTTTATAGGTCGAGTAGTATAGCCAGGTT Il24-H3L: AGGTCAGGAATACTTACGTCGTTATGGTCACTAATGGGAAGCATGGA Il24-H3R: AAAACCGCTGGTGTGCACTCTTATAGGTCGAGTAGTATAGCCAGGTT Krt14-H1L: AGGTCAGGAATACTTACGTCGTTATGGTGGCGGTTGGTGGAGGTCAC Krt14-H1R: CCATGACCTTGGTGCGGATCTTATAGGTCGAGTAGTATAGCCAGGTT Krt14-H2L: AGGTCAGGAATACTTACGTCGTTATGGAAAGAGTGAAGCCTATAGGG Krt14-H2R: AGGAAGGACAAGGGTCAAGTTTATAGGTCGAGTAGTATAGCCAGGTT

#### Evolutionary analysis of cytokines/receptors

We retrieved the protein family containing IL24 from Pfam and ECOD databases (*68, 69*). Pfam classifies proteins using sequences while ECOD takes similarity in protein structure into consideration. IL24 belongs to the Pfam family IL10 (PF00726), which is a member of the Pfam clan 4H cytokine (CL0053). 4H cytokine clan is equivalent to the 4-helical cytokine homologous group of ECOD, and we included all the 29 Pfam families from this clan in our study. We identified Pfam domains in each human protein from Uniprot using HMMER (e-value < 0.00001) (*70, 71*). A total of 59 human proteins contained Pfam domains from the 4H cytokine clan, and we extracted the sequences of these domains and aligned them using PROMALS3D (*72*) (Table S2). The multiple sequence alignment (MSA) of these Pfam domains were used for phylogenetic analysis by RAxML (-m PROTGAMMAAUTO) (*73*). After initial alignment, we picked representative cytokine from each clade that highlighted in yellow from Table S2, and used the same method to generate a smaller phylogenic tree for presentation.

We identified the receptors for all human cytokines based on literature (Table S2). We identified Pfam domains in these cytokine receptors using HMMER and found that majority (35 out of 40) of them contain >=2 tandem immunoglobulin-like (Ig-like) domains in their extracellular regions. We built MSA for two Ig-like domains from these receptors using the following approach. First, we focused on receptors containing two Ig-like domains and obtained the MSA of the tandem Ig-like domains in these receptors. Second, for each cytokine receptor with >= 3 Ig-like domains, we iterated all combinations of two Ig-like domains from it and identified the combination showing maximal sequence similarity, measured by BLOSUM55 matrix to the MSA we built in the first stage. We extracted regions for the best combination for each receptor and concatenated the sequences for the two Ig-like domains to represent this receptor. Finally, we aligned the sequences of two representative Ig-like domains from all the receptors with >= 2 such domains using PROMALS3D, and the resulting MSA was used to reconstruct the phylogeny of these receptors through RAxML.

#### Germ-free mice wounding

Germ-free (GF) C57BL/6 wild-type (WT) mice were kept in germ-free flexible film isolators (Class Biologically Clean Ltd) at Rockefeller University. For wounding experiments, GF C57BL/6 mice were exported to isocages bioexclusion system (Tecniplast, PA, USA) and housed in isocages for the duration of the experiment. Wounding of GF mice was performed in a sterile hood using sterile autoclaved instruments. Wounding of specific-pathogen-free (SPF) C57BL/6 WT mice was performed in the same hood after GF mice were transferred into the isocages. Both GF and SPF mice were then housed in the isocages under the same conditions for 1 or 5 days as described before harvesting skin wounds. Mice housed in the isocages were provided with autoclaved food and water.

#### Fluorescence-activated cell sorting

In order to isolate and stain EpdSCs from the homeostatic mouse back skin, subcutaneous fat was removed from the skin with a scalpel, and the skin was placed dermis side down on 0.25% trypsin (Gibco) and 0.1 mg/ml DNase at 37 °C for 45 minutes while shaking gently. For isolating Day-1 wound edge EpdSCs, skin wounds were first excised at about 0.5 mm from the wound edge. Subcutaneous fat was then removed, and the skin was placed on a Whatman filter paper, faced down to be soaked entirely in trypsin, and incubated for 18 minutes while shaking gently. For Day-5 or 7 wounds, wounds were excised at 0.5 mm from the wound edge, placed on a Whatman filter paper, faced down to be soaked entirely in 50 mM EDTA in PBS, and incubated at 37°C for 1 hour while shaking gently. After the incubation, the wound edge epidermis including the migrating tongue was carefully dissected and isolated from the dermis under a dissection microscope. The isolated epidermis was then incubated in trypsin for 12 minutes while shaking gently. Single-cell suspensions were obtained by scraping the skin to remove the epidermis and hair follicles from the dermis of homeostatic skin or Day-1 wounds. Single-cell suspensions for Day-5 or 6 wounds were obtained by pipetting the suspension to release single cells. Cells were then filtered through 70 µm, followed by 40 µm strainers. Cell suspensions were incubated with the indicated antibodies for 30 minutes on ice. The following anti-mouse antibodies were used for FACS: α6-integrin-PE or BV650 (BD Pharmingen, 1:1,000), CD34-efluro660 or BV421 (eBiosciences, 1:100), Sca-1-PerCP-Cy5.5 (Biolegend, 1:1,000), CD45-APC-Cy7 (Biolegend,1:200), CD31-PE-Cy7 (Biolegend,1:300), biotin-CD117 (Biolegend, 1:200), CD140a-APC (Biolgened, 1:100), Streptavidin-PE-Cy7 (eBioscience) 1:500, CD90-BV421 (Biolegend, 1:200). For biotin-conjugated primary antibodies, after washing with FACS buffer, cells were incubated with Streptavidin PE-Cy7 (1:500). DAPI was used to exclude dead cells. Cell isolations were performed on FACS Aria sorters running FACS Diva software (BD Biosciences). FACS analyses were performed using LSRII FACS Analyzers, and results were analyzed with FlowJo software.

For the analysis of dermal cells at the wound site, wound tissue was isolated from the back skin, keeping margins as close as 0.5-1 mm. Tissue was minced in media (RPMI with L-glutamine, sodium pyruvate, acid-free HEPES, penicillin and streptomycin), added with Liberase TL (Roche; 25 μg/ml), and digested for 60-90 minutes at 37°C while shaking gently. The digest reaction was stopped by adding 20 ml of 0.5 M EDTA and 1 ml of 10% DNase solution. Cells were filtered through a 70 µm strainer and stained with the following antibodies from Biolegend: α6-integrin-PE (1:1,000), CD45-APC-Cy7 (1:200), CD31-PE-Cy7 (1:300), CD11b-BV421 (1:1,500), MHCII-AF700 (1:1,000), CD45-APC-Cy7 (1:200), CD140a-APC (1:100), ITGA5-Ax488 or APC (1:100), Ly6G-PE or APC (1:500). Dead cells were excluded using a LIVE/DEAD Fixable Blue Dead Cell Stain Kit (Molecular Probes). FACS analyses were performed using LSRII FACS Analyzers and results were analyzed with FlowJo software.

#### Bulk RNA-seq and quantitative RT-PCR

Total RNA from sorted EpdSCs, endothelial cells, dermal fibroblasts, and innate immune cells was purified using the Direct-zol RNA MiniPrep kit (Zymo Research) per the manufacturer’s instructions. DNase treatment was performed to remove genomic DNA (RNase-Free DNase Set, Qiagen). The quality of RNA samples was determined using an Agilent 2100 Bioanalyzer, and all samples for sequencing had RNA integrity (RIN) numbers >8. cDNA library construction using the Illumina TrueSeq mRNA sample preparation kit was performed by the Weill Cornell Medical College Genomic Core facility (New York, NY), and cDNA libraries were sequenced on an Illumina HiSeq 2000 or Illumina Novaseq 6000 instruments.

The bulk RNA-seq data analysis was mainly processed in R (version 4.0) environment. The reference genome sequence was fetched from BSGenome.Mmusculus.UCSC.mm10 package (https://bioconductor.org/packages/release/data/annotation/html/BSgenome.Mmusculus.UCSC.mm10.html); the GTF file was fetched from TxDb.Mmusculus.UCSC.mm10.knownGene package (https://bioconductor.org/packages/release/data/annotation/html/TxDb.Mmusculus.UCSC.mm10.knownGene.html). The fastq files were aligned to reference genome by Salmon (version 1.4.0, https://salmon.readthedocs.io/en/latest/salmon.html), and the counts for each feature were calculated by Salmon. The counting results were imported into DESeq2 object by tximport (https://bioconductor.org/packages/release/bioc/html/tximport.html). For real-time PCR, equivalent amounts of RNA from FACS-purified cells were reverse-transcribed using the SuperScript™ VILO™ cDNA Synthesis Kit (ThermoFisher Scientific). cDNAs were normalized to equal amounts using primers against *Eef1a1, Ppib*, or *Hprt*. cDNAs were mixed with indicated gene-specific primers and SYBR green PCR Master Mix (Sigma), and qRT-PCR was performed on an Applied Biosystems 7900HT Fast Real-Time PCR system.

#### 10x single-cell RNA-seq analysis

The raw fastq files of 10X data were mapped to mouse genome (mm10), and the gene expression of each gene in each cell was estimated by the count function of Cell Ranger (v 3.0.2). The counting matrices of the two samples were then merged by the aggr function of Cell Ranger. The “.cloupe” file was applied for data visualization with Loupe Browser (v 3.0.0).

More customized analyses were processed by Seurat (v 3.0.0) which was developed on R language (version 3.5.2). The following steps were derived from Seurat vignette. First, the filtered counting matrices of the samples were loaded into Seurat object. The features detected in less than five cells were removed. The proportion of mitochondrial genes oriented UMI counts (percent.mt) was also estimated. Then, the Seurat object was subjected to log normalization (Seurat::NormalizeData) and variable features identification (Seurat::FindVariableFeatures). After this step, amount 2000 variable features were identified by vst method. To merge the Seurat objects for all samples, the CCA-based workflow was applied. After merging all samples, the cells with the following criteria were removed: (i) too few genes detected (nFeature_RNA < 200); (ii) potential doublets (nCount_RNA > 99% quantile of UMI counts); (iii) potential cell debris (percent.mt > 10%). After removing low quality cells, a principal component analysis was performed (Seurat::RunPCA). The PCs used was determined by an Elbow plot (Seurat::ElbowPlot). In this case, we decided to use the first 15 PCs for the following steps, including identify neighbors (Seurat::FindNeighbors), made UMAP projection (Seurat::RunUMAP). Finally, the clusters were identified by using Louvain clustering with resolution as 0.5 (Seurat::FindClusters). The UMAP projection and clustering information were extracted and imported into Loupe Browser for more customized visualization.

#### EdU and pimonidazole injections

In order to label mitotic cells with EdU, mice were injected intraperitoneally with thymidine analogue 5-Ethynyl-2′-deoxyuridine (EdU, 50 μg/g) (Sigma-Aldrich) 3 hours before sample collection. For labeling tissue hypoxia, pimonidazole (Hypoxiaprobe) was prepared as 100 mg/ml in 0.9% saline, and was injected intraperitoneally (60 mg/kg) 1.5 to 2 hours before sample collection.

#### Tamoxifen treatment on mice

Mice expressing *Krt14-CreER*, as well as their wild-type controls, were treated with the topical application of 0.1% 4-Hydroxytamoxifen (4OH-Tam) diluted in 100% ethanol for 4 days, to manipulate the gene expression in the epidermis. After three days of resting period, the experiments were performed on the back skin of mice as indicated.

#### Doxycycline treatment on mice

Second telogen mice expressing *Krt14-rtTA*, as well as their control littermates, were put on a high-dose doxycycline (Dox, 2 mg/kg) food chow starting 2 days before the first punch biopsy. The mice were also injected intraperitoneally with 25 μg of Dox per gram of body weight at the time of first punch biopsy. For neonatal mice experiments, pregnant females were put on the high-dose Dox chow one day before they gave birth. Neonatal mice skins were harvested 48, 72, 96 hours after the start of doxy chow.

#### In utero lentiviral transduction

Concentrated lentiviral solutions were produced, and ultrasound-guided *in utero* injection of concentrated lentivirus was performed in the Comparative Biology Center at The Rockefeller University as described previously (*46*). Specifically, female mice were anesthetized with isoflurane at embryonic day 9.5, and 500 nL to 1 µL of lentivirus was injected into the amniotic sacs of the animal to selectively transduce individual progenitors within the surface ectoderm that will give rise to the skin epithelium.

#### Histology

Mouse back skin was dissected, and fixed with 4% paraformaldehyde diluted in PBS at 4°C overnight. After extensive washing with PBS, the tissues were sent to Histowiz for processing as well as H&E and Trichrome staining.

#### Toluidine blue staining and TEM

Skin samples were fixed in 2% glutaraldehyde, 4% paraformaldehyde, and 2 mM CaCl_2_ in 0.1 M sodium cacodylate buffer (pH 7.2) for >1 hour at room temperature, post-fixed in 1% osmium tetroxide, and processed for Epon embedding; ultrathin sections (60–65 nm) were counterstained with uranyl acetate and lead citrate. Images were acquired with a transmission electron microscope (TEM, Tecnai G2-12; FEI, Hillsboro, OR) equipped with a digital camera (AMT BioSprint29). Semithin sections (800 nm) were stained with toluidine blue and photographed with a Zeiss Axio Scope equipped with a Nikon Digital Sight camera.

#### IL24 receptor reconstitution and Hif1α KO

For IL24 receptor reconstitution, either a GFP control or a mouse cDNA encoding IL20RB was cloned into pTY-EF1A-puroR-2a lentiviral vector, and either a GFP control or a mouse cDNA encoding IL22RA1 and IL20RA were cloned into pTY-EF1A-HygromycinR-2a lentiviral vector. Lentivirus was packaged in 293TN cells and then used to infect wild-type or *Krt14*CreER^+^;*Hif1αfl/fl* keratinocytes, which were selected by puromycin 1 µg/ml and hygromycin 50 µg/ml for a week. For floxing out *Hif1α* exon2 in *Krt14*CreER^+^;*Hif1αfl*/fl cells to generate *Hif1*α loss of function cells, 3 μM 4-Hydroxytamoxifen (4OH-Tam) was added to the culture for 4 days. Alternatively, guide RNA targeting *Hif1α (74*) was cloned into pLentiCRISPRv2-blasticidin construct (Addgene Plasmid #98293). Lentivirus was packaged in 293TN cells and then used to infect GFP control or IL24 receptor reconstituted keratinocytes, which were selected by blasticidin (3 μg/ml, InvivoGen) for 4 days prior to the experiments.

#### Immunoblot analysis

Cells were lysed in chilled 1x RIPA buffer (10x stock, EMD Millipore) diluted in PBS containing 1 tablet of cOmplete EDTA free protease inhibitor and PhosSTOP phosphatase inhibitor for 30 minutes on ice. Protein was quantified using a Pierce BCA protein quantification kit. 20 μg of total protein lysates were loaded and separated on NuPAGE 4–12% Bis-Tris gels (Thermo Scientific). Proteins were transferred to nitrocellulose membranes, blocked for 1 hour with 5% milk in TBS-T, and incubated with the indicated primary antibodies diluted in TBS-T at 4°C overnight. Membranes were washed in TBS-T and incubated in HRP-coupled secondary antibodies at room temperature. Proteins were detected by chemiluminescence using ECL (Thermo Scientific) in a Bio-Rad ChemiDoc Imager. The following primary antibodies and dilutions were used: vinculin (Sigma, V9131 1:2000), HIF-1α (Cayman Chemical, 10006421, 1:1000), STAT3 (124H6, Cell Signaling, 1:1000), p-STAT3 (D3A7, Cell Signaling 1:1000), LDHA (21799-1-AP, Proteintech Group, 1:5000). Western blot images were processed using Adobe Photoshop CS5.

#### ATAC-Seq library preparation and sequencing

ATAC-seq was performed on 70,000 FACS-purified cells from control and Day-1 wounded samples and processed as previously described (*48*). Briefly, cells were lysed in ATAC lysis buffer for 5 minutes and then transposed with TN5 transposase (Illumina) for 30 minutes at 37°C. Samples were uniquely barcoded, and the sequencing library was prepared according to manufacturer guidelines (Illumina). Libraries were sequenced on Illumina NextSeq 500. 40-bp paired-end ATAC-seq FASTQs were aligned to the mm10 genome from the Bsgenome.Mmusculus.UCSC.mm10 Bioconductor package (version 1.4.0) using Rsubread’s align method in paired-end mode with fragments between 1 to 5000 base-pairs considered properly paired (*75*). Normalized, fragment signal bigWigs were created (*76*). Peak calls for each replicate were made with MACS2 software in BAMPE mode (*77, 78*).

#### Cut and Run-Seq analysis

Cultured EpdSCs from GFPctrl_21%O2 (24hr) and IL22RA/IL20RB_1%O2 (24 hr) were trypsinized into single cell suspensions, and CUT&RUN was performed as previously described with minor modifications indicated below (*49*). Briefly, 650,000 cells were resuspended in crosslinking buffer (10 mM HEPES-NaOH pH 7.5, 100 mM NaCl, 1 mM EGTA, 1 mM EDTA, 1% formaldehyde) and rotated at room temperature for 10 minutes. Crosslinked cells were quenched with glycine at a final concentration of 0.125 M for 5 minutes at room temperature. Cells were washed with cold PBS and resuspended in NE1 buffer (20 mM HEPES-KOH pH7.9, 10 mM KCl, 1mM MgCl2, 1 mM DTT, 0.1% triton X-100 supplemented with Roche complete protease inhibitor EDTA-free) and rotated for 10 minutes at 4°C. Nuclei were washed twice with CUT&RUN wash buffer (20 mM HEPES pH7.5, 150 mM NaCl, 0.5% BSA, 0.5 mM spermidine supplemented with protease inhibitor) and incubated with concanavalin-A (ConA) beads washed with CUT&RUN binding buffer (20 mM HEPES-KOH pH 7.9, 10 mM KCl, 1 mM CaCl_2_, 1 mM MnCl_2_) for 10 minutes at 4°C. ConA-bead-bound nuclei were incubated CUT&RUN antibody buffer (CUT&RUN wash buffer supplemented with 0.1% triton X-100 and 2 mM EDTA) and antibody at 4°C overnight. After antibody incubation, ConA-bead-bound nuclei were washed once with CUT&RUN triton wash buffer (CUT&RUN wash buffer supplemented with 0.1% triton X-100) then resuspended and incubated at 4°C for 1 hour in CUT&RUN antibody buffer and 2.5 μL pAG-MNase (EpiCypher). ConA-bound-nuclei were then washed twice with CUT&RUN triton wash buffer, resuspended in 100μL of triton wash buffer, and incubated on ice for 5 minutes. Each 100 µl ConA-bound-nuclei was added with 2 µL 100 mM CaCl_2,_ mixed gently, and incubated on ice for 30 minutes. After adding 100 μL 2x stop buffer (30 mM EGTA), the reaction was incubated at 37°C for 10 minutes. After incubation, ConA-bound-nuclei were captured using a magnet, and the supernatant containing CUT&RUN DNA fragments was collected. The supernatant was incubated at 70°C for 2 hours with 2 μL 10% SDS and 2.5 μL 20mg/mL proteinase K. DNA was purified using PCI and overnight ethanol precipitation with glycogen at -20°C, and was resuspended in 15 μL of buffer EB. CUT&RUN sequencing libraries were generated using NEBNext Ultra II DNA Library Prep Kit for Illumina and NEBNext Multiplex Oligos for Illumina (Index Primer Set 1 and 2). PCR-amplified libraries were purified using 1.2x ratio of AMPure XP beads and eluted in 15 μL 0.1x TE buffer. All CUT&RUN libraries were sequenced on Illumina NextSeq using 40-bp paired-end reads. Reads were aligned to reference genome (mm10) using Bowtie2 (version 2.2.9) and deduplicated with Java (version 2.3.0) Picard tools (http://broadinstitute.github.io/picard). Reads were flittered to reads smaller or equal to 120 bp using samtools (version 1.3.1). BAM files for each replicate were combined using samtools. Bigwigs were generated using deeptools (version 3.1.2) with RPKM normalization and presented by Integrative Genomics Viewer (IGV) software. Peaks were called using SEACR using a stringent setting and a numeric threshold of 0.01. Peaks were further filtered to have peaks scores greater than 600 for a set of high confident peaks per antibody and condition. The motif analysis was performed with HOMER (version 4.10).

### QUANTIFICATION AND STATISTICAL ANALYSIS

Group sizes were determined on the basis of the results of the preliminary experiment and mice were assigned at random to groups. The number of animals shown in each figure is indicated in the legends as *n =* x mice per group and in times, and data are presented with multiple measurements per animal. Experiments were not performed in a blinded fashion. Statistical analysis was calculated using Prism software (GraphPad). All error bars are mean ± SEM. Experiments were independently replicated, and representative data are shown. Paired two-tailed Student’s t-tests were used to ascertain statistical significance between two groups, and one-way ANOVA was used to assess statistical significance between three or more groups with one experimental parameter; *; p < 0.05, **; p< 0.01, ***; p < 0.001, ****; p < 0.0001, ns; not significant. See figure legends for more information on statistical tests.

