## Supplemental figures and tables for "A tissue injury repair pathway distinct but parallel to host pathogen defense"

### SUPPLEMENTAL FIGURES AND LEGENDS

**Figure S1 IL24 is specifically produced by epithelial stem cells near the wound site**

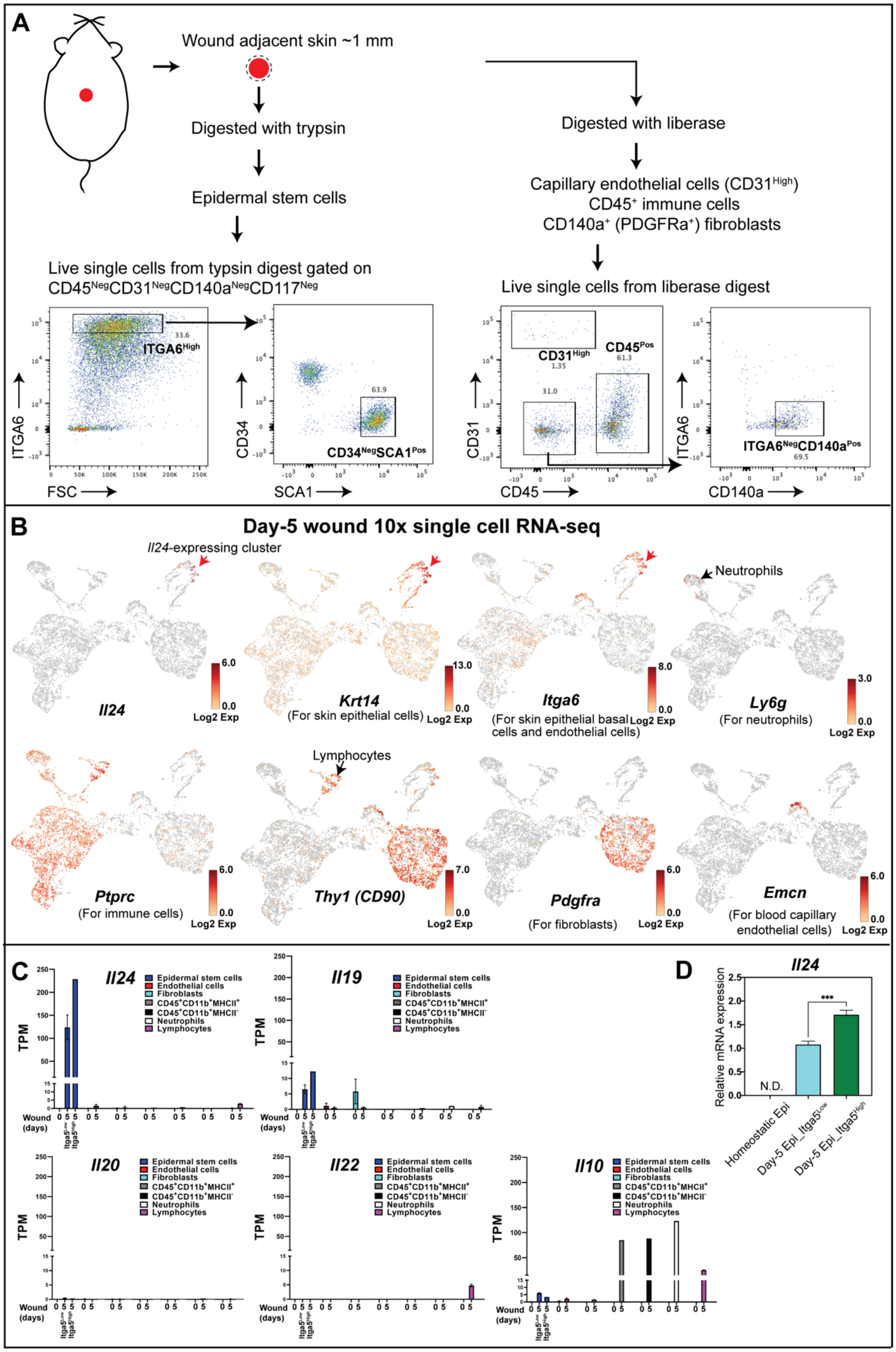

**Supplemental Figure 1**

(A) Schematic and flow panels for purification of different cell populations from the wound edge by fluorescence activated cell sorting (FACS).

(B) 10x single cell RNA-seq results from the cells consisting of Day-5 wounds. Red arrows denote the *Il24*-expressing cell clusters, highly positive for EpdSC markers *Krt14* and *Itga6*.

(C) IL10 cytokine family expression from RNA-seq performed on FACS-purified cell populations isolated from homeostatic skin (Day-0) and Day-5 wounds. EpdSCs isolated from Day-5 wounds were further separated based on the expression level of integrin-α5 (i.e. Itgα$5^{low}$ vs. Itgα$5^{High}$), a marker of the migrating epidermal tongue. Note that only IL24 and to a lesser extent IL19 are uniquely activated by EpdSCs following injury.*

(D) q-RT-PCR analysis of *Il24* mRNA expression in $SCA1^{+}$ EpdSCs that were FACS-purified from homeostatic skin and Day-5 wounds. EpdSCs isolated from Day-5 wounds were further separated based on the expression level of integrin-α5 (i.e. Itgα$5^{low}$ vs. Itgα$5^{High}$), a marker of the migrating epidermal tongue. [n=5 mice]*

*The data shown in (C) and (D) are presented as mean ± SEM. Statistical significance was determined using Student's t-tests; ***; $p < 0.001$ and N.D: not detected.

**Figure S2 Loss of IL24 reduced p-STAT3 and epithelial proliferation at the wound edge**

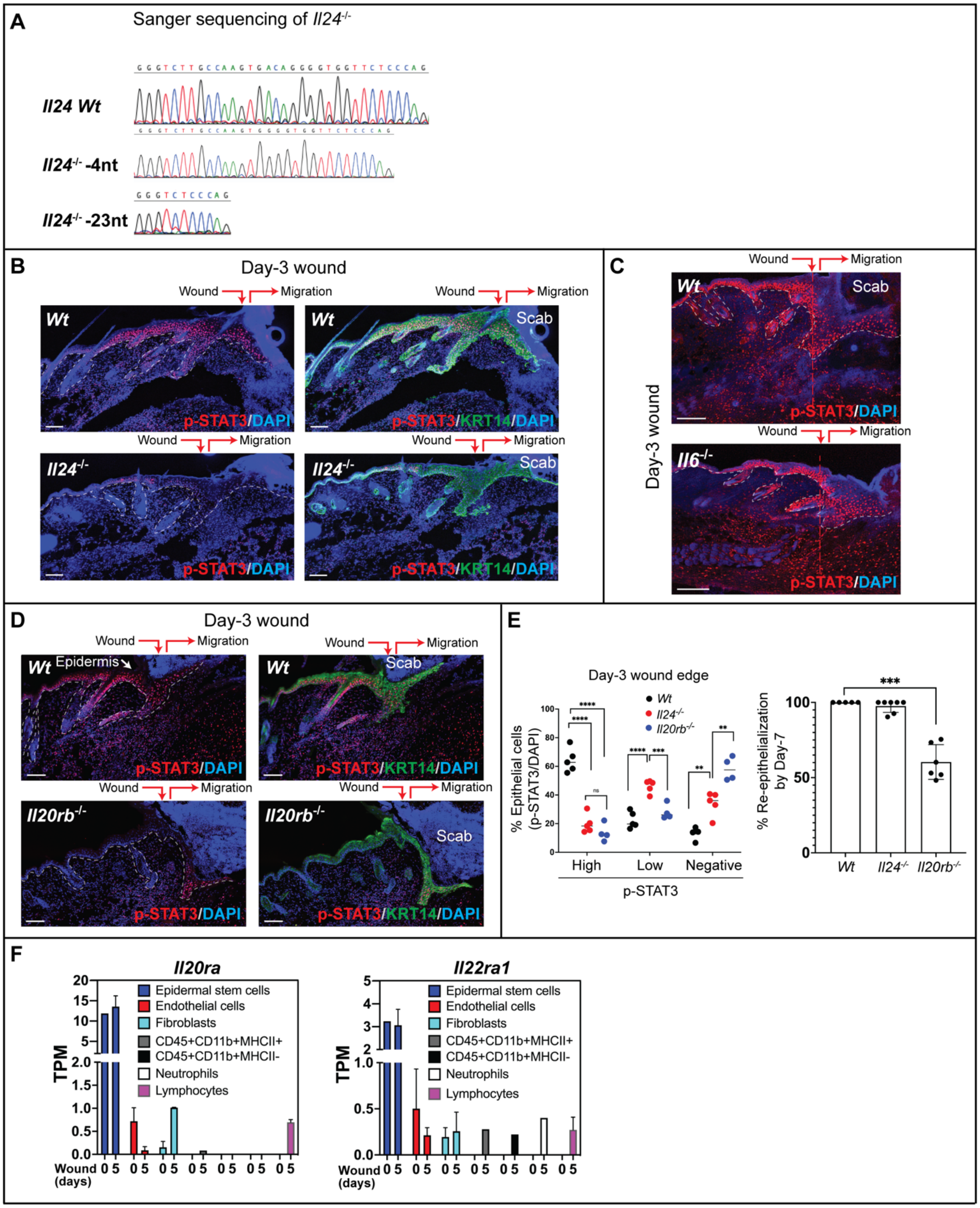


#### Supplemental Figure 2

(A) Sanger sequencing of the *Il24* exon2 locus near the guide RNA-binding site in *Il24 Wt* or *Il24*$^{-/-}$ mutants.

(B) Fluorescence microscopy of sagittal sections of Day-3 wounds from *Wt* and *Il24*$^{-/-}$ mice immunolabeled for p-STAT3 and KRT14 and stained with DAPI to label nuclei. Wound site and direction of epidermal migration are indicated. Scale bar: 100 µm. [n=5 mice for each genotype]

(C) Fluorescence microscopy of sagittal sections of Day-3 wounds from *Wt* and *Il6*$^{-/-}$ mice immunolabeled for p-STAT3 and stained with DAPI to label nuclei. Wound site and direction of epidermal migration are indicated, along with angle of wound (denoted by red dotted line). Scale bar: 100 µm. [n=5 mice for each genotype]

(D) Fluorescence microscopy of sagittal sections of Day-3 wounds from *Wt* and *Il20rb*$^{-/-}$ mice immunolabeled for p-STAT3 and KRT14 and stained with DAPI to label nuclei. Wound site and direction of epidermal migration are indicated. Scale bar: 100 µm. [n=5 mice for each genotype]

(E) Quantifications of the percentage of epithelial cells expressing different levels of p-STAT3 (left), and the percentage of re-epithelialization 7 days post-wounding in *Wt, Il24*$^{-/-}$ and *Il20rb*$^{-/-}$ mice. [*Wt*: 5 mice, *Il24*$^{-/-}$: 5-7 mice, *Il20rb*$^{-/-}$: 4-6 mice; dots indicate data from individual mice]*

(F) *Il20ra* and *Il22ra1* expression from RNA-seq performed on FACS-purified cell populations from homeostatic skin (Day-0) and Day-5 wounds. TPM: Transcripts per kilobase million. [n=5 mice]*

*The data shown in (E) and (F) are presented as mean ± SEM. Statistical significance was determined using Student's t-tests; ****; $p < 0.0001$, ***; $p < 0.001$, and **; $p < 0.01$.

**Figure S3 Epithelial-expressed IL24 coordinates dermal repair and re-epithelialization**

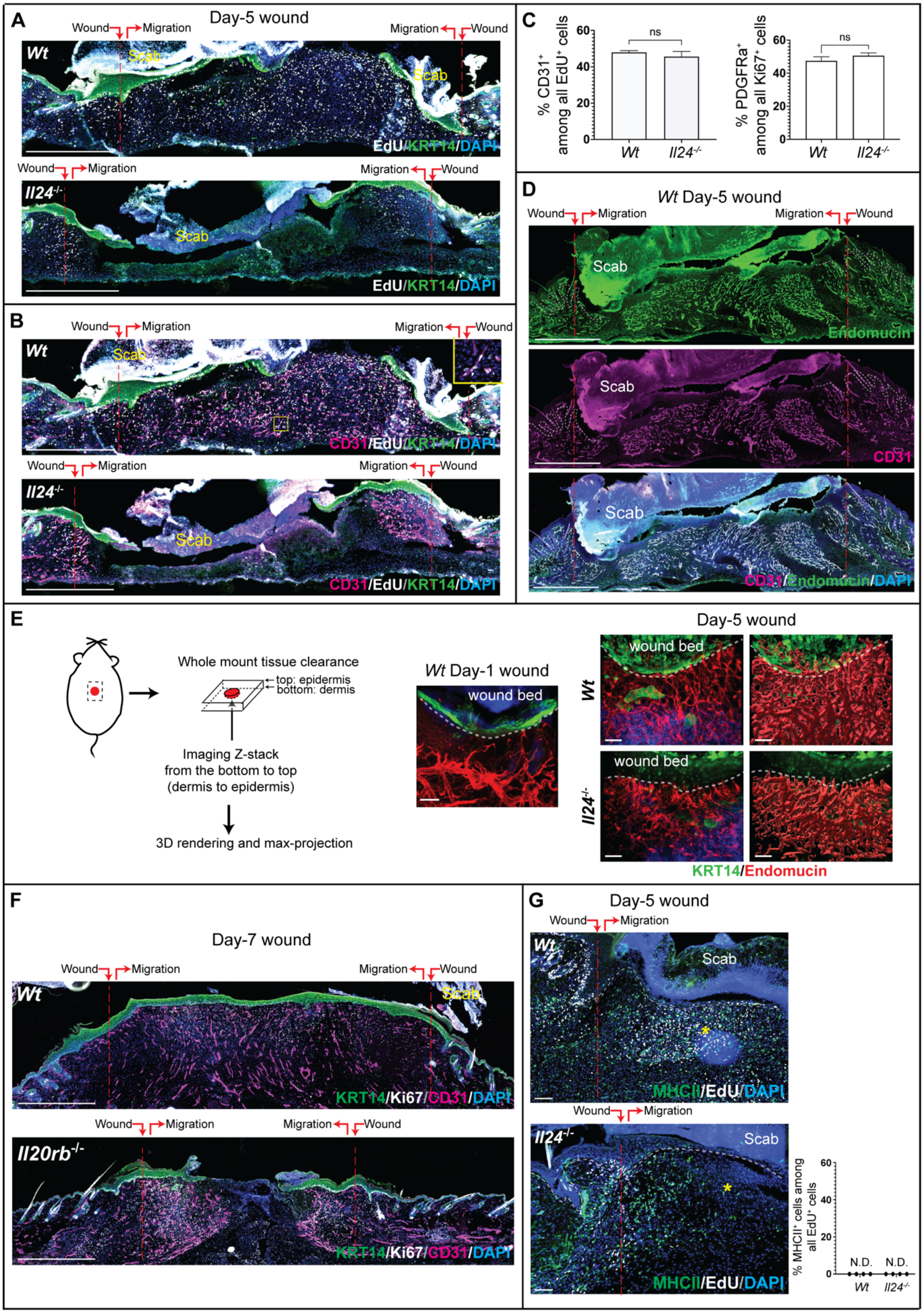

**Supplemental Figure 3**

(A) Fluorescence microscopy of sagittal sections of whole Day-5 wounds from *Wt* and *Il24*$^{-/-}$ mice immunolabeled for EdU and KRT14 and stained with DAPI to label nuclei. Some scabs above the wounds have high autofluorescence, especially in the EdU channel (Ax-488 and pseudo-colored in white). Wound site and direction of epidermal migration are indicated. Scale bar: 500 µm. [*Wt*: n=5 mice, *Il24*$^{-/-}$: n=9 mice]

(B) Fluorescence microscopy of sagittal sections of whole Day-5 wounds from *Wt* and *Il24*$^{-/-}$ mice immunolabeled for CD31, EdU and KRT14, and stained with DAPI to label nuclei. The boxed region is magnified in the top image to highlight proliferating blood capillaries (EdU$^{+}$CD31$^{+}$). Some scabs above the wounds have high autofluorescence, especially in the EdU channel (Ax-488 and pseudo-colored in white). Wound site and direction of epidermal migration are indicated. Scale bar: 500 µm. [*Wt*: n=5 mice, *Il24*$^{-/-}$: n=9 mice]

(C) Quantifications of the percentage of CD31$^{+}$ cells among all dermal mitotic (EdU$^{+}$) cells (left), and PDGFR$^{+}$ cells among all dermal proliferating (KI67$^{+}$) cells (right). [n=5 for each genotype]*

(D) Fluorescence microscopy of sagittal sections of Day-5 wounds from *Wt* mice immunolabeled for Endomucin and CD31 and stained with DAPI to label nuclei. Wound site and direction of epidermal migration are indicated. Scale bar: 500 µm. [n=3 mice]

(E) Schematic and images of whole-mount immunofluorescence microscopy and 3D image reconstruction performed on *Wt* Day-1 wound and Day-5 wounds from *Wt* and *Il24* null mice immunolabeled for Endomucin and KRT14. [n=3 mice for each genotype]

(F) Fluorescence microscopy of sagittal sections of Day-7 wounds from *Wt* and *Il20rb*$^{-/-}$ mice immunolabeled for KRT14, Ki67, and CD31, and stained with DAPI to label nuclei. Wound site and direction of epidermal migration are indicated. Scale bar: 500 µm. [*Wt*: n=3 mice, *Il20rb*$^{-/-}$: n=4 mice]

(G) Fluorescence microscopy of sagittal sections of Day-5 wounds from *Il24 Wt* and *Il24*$^{-/-}$ mice immunolabeled for MHCII (found only on professional antigen presenting cells, typically immune cells) and EdU and stained with DAPI to label nuclei. Asterisks (*) denote corresponding dermal wound regions lacking MHCII$^{+}$ cells in *Il24*$^{-/-}$ but not in *Wt* mice. Wound site and direction of epidermal migration are indicated. Scale bar: 100 µm; white dotted lines: epidermal-dermal border. The percentage of MHCII$^{+}$ cells among all dermal $EdU^+$ cells is quantified. [n=5 for each genotype]*

*The data shown in (C) and (G) are presented as mean ± SEM. Statistical significance was determined using Student's t-tests; ns; not significant and N.D.; not detected.

**Figure S4 Ultrastructural analysis of Day-5 wounds by transmission electron microscopy (TEM)**

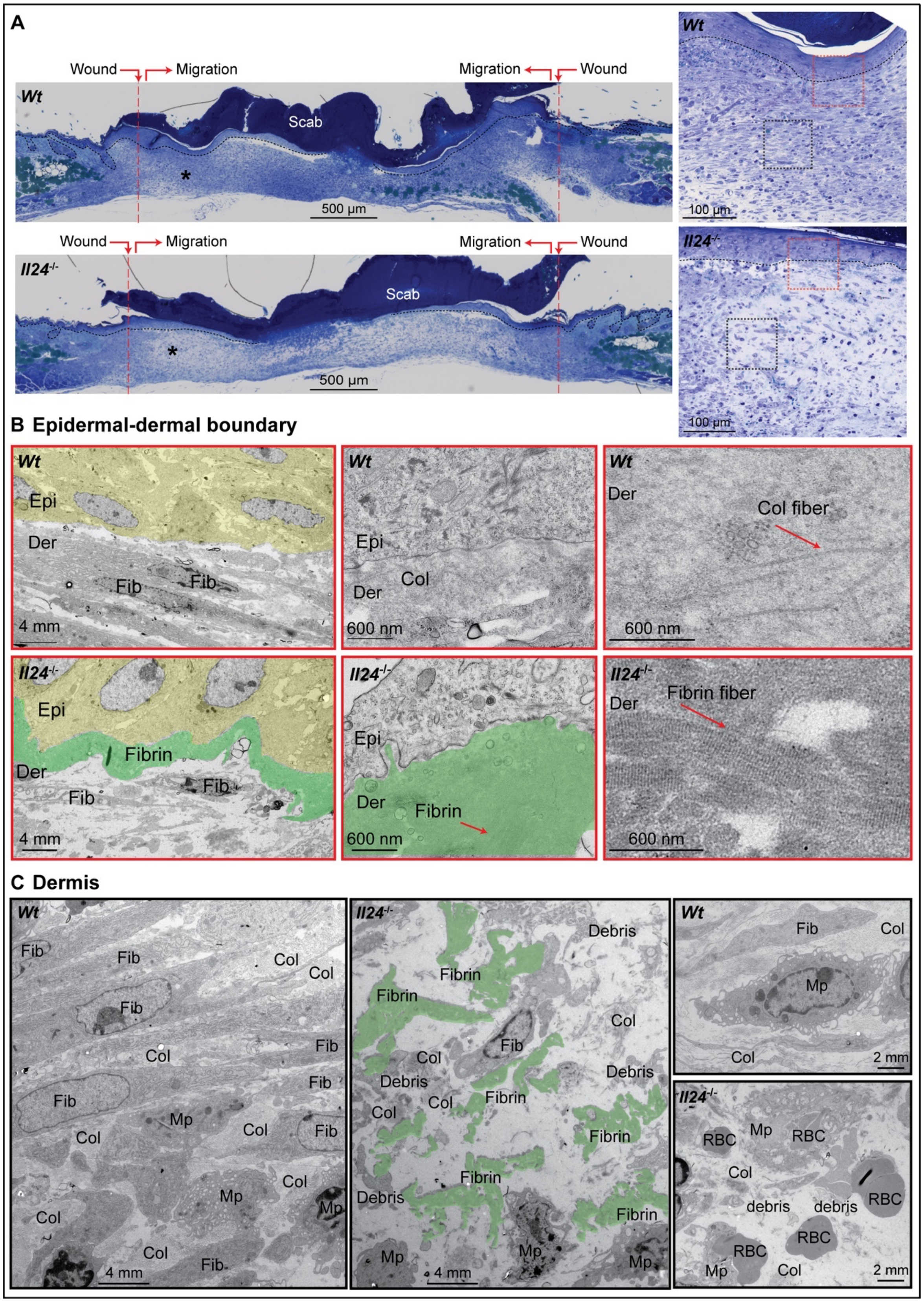

**Supplemental Figure 4**

(A) **Left**: Images of Toluidine blue staining performed on the semithin (800 nm) sections of Day-5 wounds from *Wt* and *Il24*$^{-/-}$ mice. Black asterisk (*) denotes corresponding wound regions where dermal cell number sharply decreases in *Il24*$^{-/-}$ compared to *Wt* mice. Wound site and direction of epidermal migration are indicated. Scale bar: 500 µm; black dotted lines: epidermal-dermal border. **Right**: Magnified images showing the reduced dermal cellularity in *Il24*$^{-/-}$ compared to *Wt* mice. Scale bar: 100 µm; black dotted lines: epidermal-dermal border; red box: refer to (B); black box: refer to (C). [representative images of the experiments performed on n=3 mice for each genotype]

(B) Transmission electron microscope (TEM) images in different magnifications of the regions of epidermal (Epi) and dermal (Der) boundary indicated as a red box in (A). Red arrows indicate the ultrastructure of collagen fibers in *Wt* and fibrin fibers in *Il24*$^{-/-}$ wounds. Fibrin is pseudo-colored in green. [representative images of the experiments performed on n=3 mice for each genotype]

(C) TEM images of dermal regions indicated as a black box in (A). Fibrin is pseudo-colored in green. Note that, in the bottom right panel, multiple RBCs are being engulfed by macrophages in *Il24*$^{-/-}$ wounds. Col: collagen bundles; Fib: fibroblast; Mp: macrophage; RBC: red blood cells. [representative images of the experiments performed on n=3 mice for each genotype]

**Figure S5 Tissue damage associated hypoxia and HIF1α in wounds are important for robust *Il24* expression**

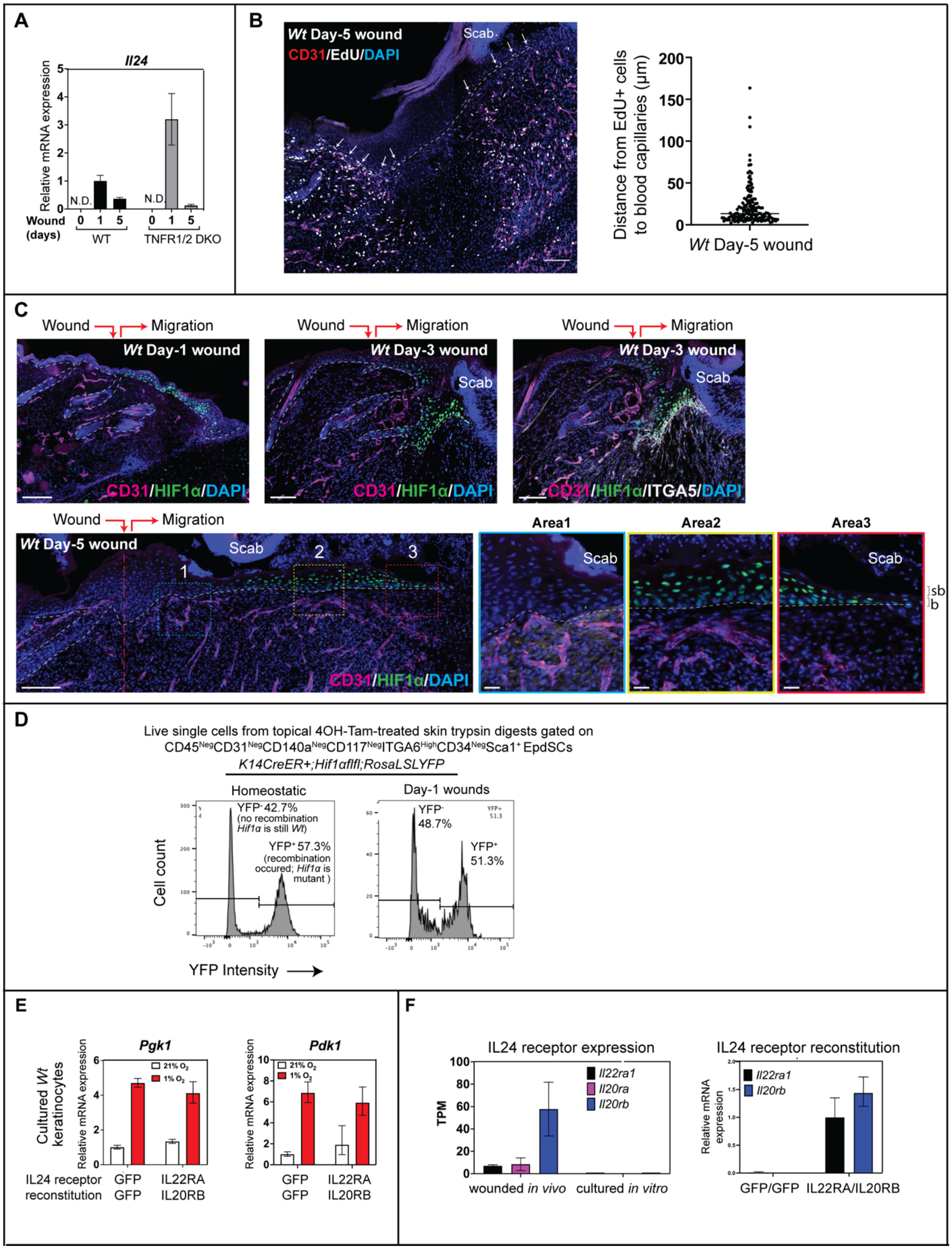

**Supplemental Figure 5**

(A) q-RT-PCR analysis of *Il24* mRNA expression in EpdSCs that were FACS-purified from homeostatic skin (Day-0), and Day-1 and 5 wounds from *Wt* and *TNRF1/2* DKO mice. *TNRF1/2* DKO mice lack TNF signaling. [n=5 mice]*

(B) Fluorescence microscopy of sagittal sections of Day-5 wound from *Wt* mice immunolabeled for CD31 and EdU and stained with DAPI to label nuclei. White arrows denote $EdU^+$ mitotic EpdSCs adjacent to blood capillaries ($CD31^+$). Wound site and direction of epidermal migration are indicated. Scale bar: 100 µm; white dotted lines: epidermal-dermal border. The distance (µm) between $EdU^+$ EpdSCs to the closest blood capillaries is quantified. [n=*3* mice]

(C) Fluorescence microscopy of sagittal sections of Day-1, 3, and 5 wounds immunolabeled for CD31, HIF1α, and Itga5, and stained with DAPI to label nuclei. Wound site and direction of epidermal migration are indicated. Images on the bottom right (Area1, 2, and 3) highlight three different regions of epidermis in Day-5 wounds (bottom left, denoted by colored boxes). Area1: wounded epidermis that has established a close association with $CD31^+$ blood capillaries, and exhibits minimal nuclear HIF1α staining; Area 2: wounded epidermis that has established a relatively close association with $CD31^+$ blood capillaries, and exhibits minimal nuclear HIF1α staining in the basal (b) EpdSCs but still shows high nuclear HIF1α in the suprabasal (sb) epidermal cells; Area 3: the tip of the epidermal migrating tongue that is still distant from $CD31^+$ blood capillaries, and exhibits relatively strong nuclear HIF1α in the basal EpdSCs. Scale bar: 100 µm (20 µm for magnified images); b: basal EpdSCs; sb: suprabasal epidermal cells; white dotted lines: epidermal-dermal border. [n=4 mice]

(D) FACS plots for purification of $YFP^+$ vs. $YFP^-$ EpdSCs from 4OH-Tam treated-*Krt14CreER*; *Hif1α*$^{fl/fl}$ mice described in Figure 5D.

(E) q-RT-PCR analyses of known HIF1α targets *Pgk1* and *Pdk1* mRNA expression in keratinocytes reconstituted with GFP (GFP/GFP) or IL24 receptors (IL22RA/IL20RB) and cultured under normoxic (21% $O_2$) or hypoxic (1% $O_2$) conditions for 24 hours. [representative results from 3 experiments]*

(F) **Left**: IL24 receptor expression from RNA-seq performed on EpdSCs FACS-purified from wound edge (wounded *in vivo*) or cultured keratinocytes (cultured *in vitro*). **Right**: q-RT-

PCR analysis of IL24 receptors *Il22ra1* and *Il20rb* in keratinocytes reconstituted with GFP (GFP/GFP) or IL24 receptors (IL22RA/IL20RB). [representative results from 3 experiments]*

*The data shown in (A), (E), and (F) are presented as mean ± SEM. N.D.; not detected.

**Figure S6 Critical roles for both hypoxia/HIF1α and STAT3 in governing robust *Il24* expression**

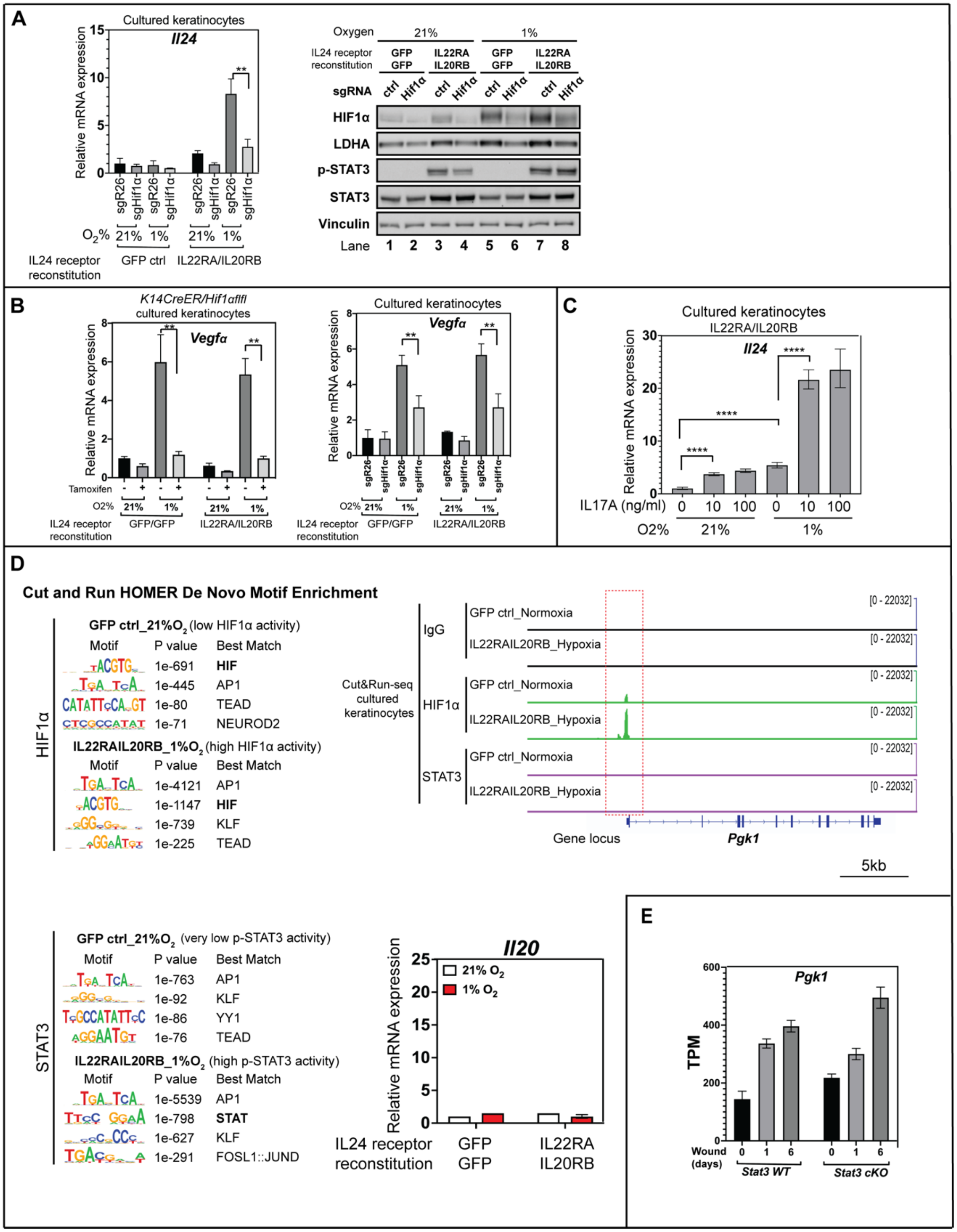

#### Supplemental Figure 6

(A) Primary epidermal keratinocytes reconstituted with either GFP (GFP ctrl) or IL24 receptor

(IL22RA/20RB) were transduced with lentivirus containing a CRISPR guide RNA to target either the *Rosa26* or *Hif1α* locus, and cultured in normoxic (21% $O_2$) or hypoxic (1% $O_2$) conditions. **Left**: q-RT-PCR analysis of *Il24* mRNA expression in the cells described. **Right**: A duplicate set of samples were immunoblotted for HIF1α, LDHA, p-STAT3, STAT3, and vinculin as the loading control. [representative of 3 independent experiments]*

(B) q-RT-PCR analyses of *Vegfa* mRNA expression in the same samples described in Figure 6B (left) and described in (A) (right).*

(C) Keratinocytes reconstituted with IL24 receptor (IL22RA/20RB) were cultured in normoxic (21% $O_2$) or hypoxic (1% $O_2$) conditions, and treated with different amounts of IL17A. q-RT-PCR analysis was then performed on the cells described to evaluate *Il24* mRNA expression [representative of 2 independent experiments]*

(D) **Left**: Unbiased motif analysis showing the enrichment of HIF1α and STAT3 binding motifs in Cut&Run samples described in Figure 6E. **Right**: Normalized peaks of HIF1α and STAT3 *in vitro* Cut&Run-seq at the *Pgk1* locus. HIF1α and STAT3 Cut&Run-seq samples were collected from *in vitro* cultured cells GFP/GFP_control_21%$O_2$_24 hrs (very low IL24 expression) and IL22RA/IL20RB_1%$O_2$_24 hrs (very high IL24 expression). q-RT-PCR analysis of *Il20* mRNA expression, a gene that is adjacent to the *Il24* locus, and to HIF1α and STAT3 binding sites, is shown in the bottom right panel.

(E) The expression of *Pgk1*, a known HIF1α target, from RNA-seq performed on EpdSCs FACS-purified from homeostatic skin (Day-0), and Day-1 and Day-6 wounds from *Wt* vs. *Stat3* cKO mice. TPM: Transcripts per kilobase million. [n=3 mice for each genotype]*

*The data shown in (A), (B), (C), and (E) are presented as mean ± SEM. Statistical significance was determined using Student's t-tests; ****; $p < 0.0001$ and **; $p < 0.01$.

#### SUPPLEMENTAL TABLES AND LEGENDS

**Table S1 A list of cytokines that activate STAT3**

| Cytokines | Corresponding Receptors | Receptor expressed in EpdSCs? |
|---|---|---|
| **IL6 family:** CLCF1, IL6, IL11, IL12, LIF, OSM | CNTFR, IL6RA/IL6ST, IL11RA/IL6ST, IL12RB1/IL12RB2, LIFR, LIFR/IL6ST,OSMR/IL31RA | Yes |
| **IL6 family:** CTF1, CNTF, GCSF, IL31, Leptin (LEP), IL27 | CLC/CNTFR, CNTFR, GCSFR/CD114, IL31RA, LEPR/IL6RA, IL27RA | No |
| **IL10 family:** IL19, IL20, IL22, IL24, IL10 | IL20RA/IL20R2, IL20R1/IL20R2 and IL22R/IL20R2, IL22R/IL10R2, IL20R1/IL20R2 and IL22R/IL20R2, IL10R1/IL10R2 | Yes |
| **IFN family:** IFNA, IFNB, IFNG | IFNAR1/IFNAR2, IFNAR1/IFNAR2, IFNGR1/IFNGR2 | Yes |
| **RTK family:** EGF, TGFA, amphiregulin (AREG), HBEGF, epiregulin (EREG), and betacellulin (BTC) | EGFR | Yes |
| **RTK family:** HGF | cMet | Yes |
| **RTK family:** PDGF, CSF-1 | PDGFR1/PDGFR2, CSF1R | No |

##### Supplemental Table 1

**A list of cytokines known to activate STAT3.** Cytokines that are known to activate STAT3 are shown and are grouped into cytokine families (first column). Corresponding cytokine receptors are shown in the second column. The cytokine receptors that have a TPM value >=1 is considered to be expressed in EpdSCs based on RNA-seq. The cytokines (highlighted in red) whose receptors are expressed in epidermal stem cells were examined for their expression in the microdissected wounded skin.

#### Table S2 IL24 protein sequence alignment across mammals

```
CLUSTAL O(1.2.4) multiple sequence alignment


House_mouse           --------------------------MSWGLQILPCLSLILLLWNQVPGLEGQEFRFGSC      34
Rat                   ------------------------MQTSLRQQILPGLSLILLVLNQVPELQGQEFRFGPC      36
Human                 -MNFQQRLQSLWTLARPFCPPLLATASQMQMVVLPCLGFTLLLWSQVSGAQGQEFHFGPC      59
Chimpanzee            WVRGPASQAASLSFSRPFCPPLLATASQMQMAVLPCLGFTLLLWSQVSGAQGQEFQFGPC      60
Black_flying_fox      ------------------------MGSPMQKAALPCLSFILLVWSWAPGVQGQEFQFGSC      36
Horse                 ------------------------MGSPMQRAALLSLSLILLLWSQRPGVQGQEFQFGSC      36
Dog                   -MN---------------ALWASSSRSTWGFLVMPCLSLILLLRSQGPGVQGQEFRFGPC      44
Pig                   ------------------------MGSPAPRAALPCLGLILLLWSQGPGVQGQEFQFGPC      36
Giant_Panda           -MVGKGR-----------ETRDQRMGFPGQTTALLCLSLILLLWSQAPGVQGQEFQFGPC      48
Polar_bear            ------------------------MGFSRQTTALLCLNLILLLWSQAPGVQGQEFQFGPC      36
Sperm_whale           ------------------------MGSPVHMSTLPCLSLILLFWSQGPGVQGQEFQFGPC      36
Killer_whale          ------------------------MGSPVRMAALPCLSLILLLWSQGPGVQGQEFQFGPC      36
Dolphin               ------------------------MGSPVRMAALPCLSLILLLWSQGPGVQGQEFQFGPC      36
                                                       :  *.: **. .     :****:** *

House_mouse           QVTGVVLPELWEAFWTVKNTVQTQDDITSIRLLKPQVLRNVSGAESCYLAHSLLKFYLNT      94
Rat                   QVTGVVLPELWEAFWTVKNTVKTQDELTSVRLLKPQVLQNVSDAESCYLAHSLLKFYLNT      96
Human                 QVKGVVPQKLWEAFWAVKDTMQAQDNITSARLLQQEVLQNVSDAESCYLVHTLLEFYLKT      119
Chimpanzee            QVKGVVPQKLWEAFWAVKDTTQAQDNITSARLLQQEVLQNVSDAESCYLVHTLLEFYLKT      120
Black_flying_fox      RVKGVDFQELWEAFQAMKDIVQAQDNITSIRLLRREVLQNVSDTESCYLIRALLKFYLNT      96
Horse                 RVEGVVLQELWAAFRAVKDVVQAQDNITGVRLLRKEVLQNVSDAESCYLSQALLKFYLDT      96
Dog                   RVQGVALRELREAFWTVKDTVQAKDNITSVRLLRKEVLQDVSDAESCYLIRALLKFYLNT      104
Pig                   RVEGIVLQELWEAFWDMKDVVQAQDNITNVRLLRKEVLQNVSEAESCYLIHSLLKFYLNT      96
Giant_Panda           RVEGVVLQELWEAFWAMKDIVQAKDNITSVRLLRKEVVQNVSGAESCYLIHALLKFYLNT      108
Polar_bear            RVEGVVLQELWEAFSAMKDIVQAKDNITSVRLLRKEVLQNVSDAESCYLIRALLKFYLTT      96
Sperm_whale           QVEGVVLQELWEAFQAMKDIAQAQDNITSVQLLRKEVLQNVSEAESCYLIHALLEFYLNT      96
Killer_whale          QVEGVVLQELWEAFQAMKDIVQAQDNITSVQLLRKEVLQNVSEAESCYLIHALLEFYLNT      96
Dolphin               QVEGVVLQELWEAFQAMKDIVQAQDNITSVQLLRKEVLQNVSEAESCYLIHALLEFYLNT      96
                      :* *:   :*  **  :*:  :::*::*. :**: :*:::** :***** ::**:*** *

House_mouse           VFKNYHSKIAKFKVLRSFSTLANNFIVIMSQLQPSKDNSMLPISESAHQRFLLFRRAFKQ      154
Rat                   VFKNYHSKIVKFKVLKSFSTLANNFLVIMSKLQPSKDNAMLPISDSARRRFLLYHRTFKQ      156
Human                 VFKNYHNRTVEVRTLKSFSTLANNFVLIVSQLQPSQENEMFSIRDSAHRRFLLFRRAFKQ      179
Chimpanzee            VFKNYHNRTVEVRTLKSFSTLANNFVLIVSQLQPSQENEMFSIRDSAHRRFLLFRRAFKQ      180
Black_flying_fox      VFKSYHEKAAEFRILKSFSTLANNFIFIASKLQPSVSRTKLGPGESARRRFLLFQRAFKQ      156
Horse                 VFKNYHGKAAEFRILKSFSTLANNFIAITSRLRPSQENEMFSISESARRRFLLFQREFKQ      156
Dog                   VFKNYLDEAADVRIRRSFSTLANNFFVIASKLQPSQEDEMFSISESARRRFLLFQRAFKQ      164
Pig                   IFKNYREKAVKFRILRSFSTLANNFVVIMSKLQPSQENEMFPISENARRRFLLFQREFKQ      156
Giant_Panda           VFKNYLDKAADSRIRKSFSTLANNFFVIVSKLQPSKENEMLSISESARRRFLLFQRAFKQ      168
Polar_bear            VFKNYLDKAADSRIRKSFSTLANNFFVIVSKLQPSQENEMFSISESARRRFLLFQRAFKQ      156
Sperm_whale           VFKNYHDKAVEFRILKSFSTLANNFIVIMSKLQPSQEKEMFSIRESARRRFLLFRRAFKQ      156
Killer_whale          VFKNYRDKAVEFGILKSFSTLANNFIVIVSKLQPSQEKEMFSISESAHRRFLLFQRAFKQ      156
Dolphin               VFKNYHDKAVEFRILKSFSTLANNFIVIVSKLQPSQEKEMFSISESARRRFLLFQRAFKQ      156
                      :**.*  . ..    :*********. * *:*:** .   :   :.*::****::* ***

House_mouse           LDTEVALVKAFGEVDILLTWMQKFYHL       181
Rat                   LDIEVALAKAFGEVDILLAWMQNFYQL       183
Human                 LDVEAALTKALGEVDILLTWMQKFYKL       206
Chimpanzee            LDVEAALTKALGEVDILLTWMQKFYKL       207
Black_flying_fox      LDIEAAQTKAFGEVDILLTWMQKFYQL       183
Horse                 LDIEAALTKAFGEVDILLTWMEKFYQP       183
Dog                   LDIQAAQTKAFGEVDILLTWMEKFYEF       191
Pig                   LDREVALTKAFGEMDILLTWMETFYQR       183
Giant_Panda           LDIQAAQTKAFGEVDILLTWMEKFYQF       195
Polar_bear            LDIQAAQTKAFGEVDILLTWMEKFYQF       183
Sperm_whale           LDREAAVTKAFGEVDILLTWMENFYH-       182
Killer_whale          LDREAAVTKAFGEVDILLTWMENFYQV       183
Dolphin               LDREAAVTKAFGEVDILLTWMEKFYQI       183
                      ** :.* .**:**:****:**:.**.
```

**Supplemental Table 2**

**IL24 protein sequence alignment across mammals.** IL24 protein sequences from different mammals were obtained from NCBI and aligned using Clustal Omega. Asterisk (*) indicates positions that have a single, fully conserved residue; Colon (:) indicates conservation between groups of strongly similar properties; Period (.) indicates conservation between groups of weakly similar properties.

**Table S3 A list of human cytokines and their receptors for homology analysis**

| Ligand | Annotation | Receptor subunit A | Receptor subunit B | Ligand | Annotation | Receptor subunit A | Receptor subunit B | Additional Receptor subunit |
|---|---|---|---|---|---|---|---|---|
| CLCF1 | Cardiotrophin-like cytokine factor 1 | CNTFR | | IFNL3 | Interferon lambda-3 | IL10RB | IFNLR1 | |
| CNTF | Ciliary neurotrophic factor | CNTFR | | IFNL4 | Interferon lambda-4 | IL10RB | IFNLR1 | |
| CSF1 | Macrophage colony-stimulating factor 1 | CSF1R | | IFNW1 | Interferon omega-1 | IFNAR1 | IFNAR2 | |
| CSF2 | Granulocyte-macrophage colony-stimulating factor | CSF2RA | CSF2RB | IL10 | Interleukin-10 | IL10RA | IL10RB | |
| CSF3 | Granulocyte colony-stimulating factor | CSF3R | | IL11 | Interleukin-11 | IL11RA | IL6ST | |
| CSH1 | Chorionic somatomammotropin hormone 1 | PRLR | | IL12A | Interleukin-12 subunit alpha | IL12RB1 | IL12RB2 | |
| CSH2 | Chorionic somatomammotropin hormone 2 | PRLR | | IL13 | Interleukin-13 | IL13RA | IL2RG | |
| CSHL1 | Chorionic somatomammotropin hormone-like 1 | PRLR | | IL15 | Interleukin-15 | IL15RA | IL2RG | |
| EPO | Erythropoietin | EPOR | | IL19 | Interleukin-19 | IL20RA | IL20RB | |
| FLT3LG | Fms-related tyrosine kinase 3 ligand | FLT3 | | IL2 | Interleukin-2 | IL2RA | IL2RB | IL2RG |
| GH1 | Somatotropin | GHR | | IL20 | Interleukin-20 | IL22RA1 | IL20RB | IL20RA |
| GH2 | Growth hormone variant | GHR | | IL21 | Interleukin-21 | IL21R | IL2RG | |
| IFNA1 | Interferon alpha-1/13 | IFNAR1 | IFNAR2 | IL22 | Interleukin-22 | IL22RA1 | IL20RB | |
| IFNA10 | Interferon alpha-10 | IFNAR1 | IFNAR2 | IL23A | Interleukin-23 subunit alpha | IL12RB1 | IL23R | |
| IFNA14 | Interferon alpha-14 | IFNAR1 | IFNAR2 | IL24 | Interleukin-24 | IL22ra1 | IL20rb | IL20RA |
| IFNA16 | Interferon alpha-16 | IFNAR1 | IFNAR2 | IL26 | Interleukin-26 | IL20ra | IL10rb | |
| IFNA17 | Interferon alpha-17 | IFNAR1 | IFNAR2 | IL3 | Interleukin-3 | IL3RA | CSF2RB | |
| IFNA2 | Interferon alpha-2 | IFNAR1 | IFNAR2 | IL34 | Interleukin-34 | CSF1R | | |
| IFNA21 | Interferon alpha-21 | IFNAR1 | IFNAR2 | IL4 | Interleukin-4 | IL4RA | IL2RG | |
| IFNA4 | Interferon alpha-4 | IFNAR1 | IFNAR2 | IL5 | Interleukin-5 | IL5RA | CSF2RB | |
| IFNA5 | Interferon alpha-5 | IFNAR1 | IFNAR2 | IL6 | Interleukin-6 | IL6RA | IL6ST | |
| IFNA6 | Interferon alpha-6 | IFNAR1 | IFNAR2 | IL7 | Interleukin-7 | IL7R | | |
| IFNA7 | Interferon alpha-7 | IFNAR1 | IFNAR2 | KITLG | Kit ligand | KIT | | |
| IFNA8 | Interferon alpha-8 | IFNAR1 | IFNAR2 | LEP | Leptin | LEPR | | |
| IFNB | Interferon beta | IFNAR1 | IFNAR2 | LIF | Leukemia inhibitory factor | LIFR | IL6ST | |
| IFNE | Interferon epsilon | IFNAR1 | IFNAR2 | OSM | Oncostatin-M | OSMR | IL31RA | |
| IFNG | Interferon gamma | IFNGR1 | IFNGR2 | PRL | Prolactin | PRLR | | |
| IFNK | Interferon kappa | IFNAR1 | IFNAR2 | THPO | Thrombopoietin | MPL | | |
| IFNL1 | Interferon lambda-1 | IL10RB | IFNLR1 | TSLP | Thymic stromal lymphopoietin | IL7R | CRLF2 | |
| IFNL2 | Interferon lambda-2 | IL10RB | IFNLR1 | | | | | |

#### Supplemental Table 3

**A list of human cytokines and their receptors for homology analysis.** A total of 59 human proteins (columns 1 and 2) that share sequence and structure homology with IL24 were bioinformatically extracted from human genome. These proteins were then subjected to initial homology alignment to generate a preliminary tree. Based on this tree, the representative cytokines from each clade/close family members highlighted in yellow, are then subjected to the same method to generate a smaller tree shown in Figure 2A. The receptors for the 59 human cytokines are listed (Columns 3-5). The receptors whose extracellular domain share homology with the IL24 receptors and contain >=2 tandem Ig-like domain were subjected to homology alignment and shown in Figure 2B.

**Table S4 A list of gene-specific q-RT-PCR primers used in the study**

| | | | | | |
|---|---|---|---|---|---|
| 1 | *mHprt_F* | GATCAGTCAACGGGGGACATAAA | 33 | *mIl20_F* | TCTACCAGACCCCTGACCAC |
| 2 | *mHprt_R* | CTTGCGCTCATCTTAGGCTTTGT | 34 | *mIl20_R* | CATTGCTTCTTCCCCACAAT |
| 3 | *mIl6_F* | TCCATCCAGTTGCCTTCTTG | 35 | *mTgfa_F* | ATCACCTGTGTGCTGATCCA |
| 4 | *mIl6_R* | GGTCTGTTGGGAGTGGTATC | 36 | *mTgfa_R* | CAAGCAGTCCTTCCCTTCAG |
| 5 | *mIfnb_F* | CCCTATGGAGATGACGGAGA | 37 | *mAreg_F* | GACTCACAGCGAGGATGACA |
| 6 | *mIfnb_R* | CTGTCTGCTGGTGGAGTTCA | 38 | *mAreg_R* | CTGTGATAACGATGCCGATG |
| 7 | *proIl1bF* | GGGCCTCAAAGGAAAGAATC | 39 | *mEreg_F* | TCTGACATGGACGGCTACTG |
| 8 | *proIl1bR* | TACCAGTTGGGGAACTCTGC | 40 | *mEreg_R* | CGCAACGTATTCTTTGCTCA |
| 9 | *mIfnaF* | ATTTTGGATTCCCCTTGGAG | 41 | *mBtc_F* | GCACAGGTACCACCCCTAGA |
| 10 | *mIfnaR* | TATGTCCTCACAGCCAGCAG | 42 | *mBtc_R* | GCCCCAAAGTAGCCTTTCTC |
| 11 | *mIl24_F* | GCCCAGTAAGGACAATTCCA | 43 | *mIl22ra1F* | AGG TCC ATT CAG ATG CTGGT |
| 12 | *mIl24_R* | ATTTCTGCATCCAGGTCAGG | 44 | *mIl22ra1R* | TAG GTG TGG TTG ACG TGGAG |
| 13 | *mIl22_F* | CCGAGGAGTCAGTGCTAAGG | 45 | *mIl20Ra_F* | GAAGAACGTGGTCCCAGTGT |
| 14 | *mIl22_R* | CATGTAGGGCTGGAACCTGT | 46 | *mIl20Ra_R* | AAGTAGCCAATTGCGGAGAA |
| 15 | *mEgf_F* | GGGAAAATGTGTCTCCCTCA | 47 | *Eef1a1_F* | AACCACCGCTAATTCAAAGCAA |
| 16 | *mEgf_R* | TCATGCCTGACACCATGATT | 48 | *E ef1a1_R* | AGGAGCCCTTTCCCATCTCAG |
| 17 | *mHbegf_F* | CAGGACTTGGAAGGGACAGA | 49 | *mIl20rb_F* | CCTCCCAGACACCTTGAAAA |
| 18 | *mHbegf_R* | CCGTGGATGCAGTAGTCCTT | 50 | *mIl20rb_R* | CAAAGAGATGCTCCGAGGAC |
| 19 | *mIl11_F* | CATTGGGATCTTTGCAGCTT | 51 | *mPpib_F* | TGATCCAGGGTGGAGACTTC |
| 20 | *mIl11_R* | GAGCTGTAAACGGCGGAGTA | 52 | *mPpib_R* | ATTGGTGTCTTTGCCTGCAT |
| 21 | *mLif_F* | AGAAGGTCCTGAACCCCACT | 53 | *mIfng_F* | GCGTCATTGAATCACACCTG |
| 22 | *mLif_R* | CCACACGGTACTTGTTGCAC | 54 | *mIfng_R* | TGAGCTCATTGAATGCTTGG |
| 23 | *mClcf1_F* | CGAGCCTGACTTCAATCCTC | 55 | *mPdk1_F* | GTGCCCCTGGCTGGGTTTGG |
| 24 | *mClcf1_R* | TACGTCGGAGTTCAGCTGTG | 56 | *mPdk1_R* | CCAGGCGTCCCATGTGCGTT |
| 25 | *mHgf_F* | ATGGGGAATGAGAAATGCAG | 57 | *mVegfa_F* | GGAGAGCAGAAGTCCCATGA |
| 26 | *mHgf_R* | CTCCCTCACATGGTCCTGAT | 58 | *mVegfa_R* | ACTCCAGGGCTTCATCGTTA |
| 27 | *mIl12a_F* | CATCGATGAGCTGATGCAGT | 59 | *mPgk1_F* | ATTCTGCTTGGACAATGGAGC |
| 28 | *mIl12a_R* | CAGATAGCCCATCACCCTGT | 60 | *mPgk1_R* | AGGCATGGGAACACCATCA |
| 29 | *mOsm_F* | TCAGGGGTCTGATGACACAA | 61 | *mIl24_3UTRF1* | GTTGTTGGCTCAGGCTTTTC |
| 30 | *mOsm_R* | GTGTGAGGTCACCCAGAGGT | 62 | *mIl24_3UTRR1* | GTTTCCAGGGAAGGTGACAA |
| 31 | *mIl19_F* | ATCCTGTCCCTGGAGAACCT | 63 | *mIl24_456F* | CACTCTGGCCAACAACTTCA |
| 32 | *mIl19_R* | AAAGAGTTGGCAATGCTGCT | 64 | *mIl24_608R* | GCTTTCACCAAAGCGACTTC |

#### Supplemental Table 4

#### A list of gene-specific qPCR primers used in the study.
